# Giant African snail genomes provide insights into molluscan whole-genome duplication and aquatic-terrestrial transition

**DOI:** 10.1101/2020.02.02.930693

**Authors:** Conghui Liu, Yuwei Ren, Zaiyuan Li, Qi Hu, Lijuan Yin, Xi Qiao, Yan Zhang, Longsheng Xing, Yu Xi, Fan Jiang, Sen Wang, Cong Huang, Bo Liu, Hengchao Wang, Hangwei Liu, Fanghao Wan, Wanqiang Qian, Wei Fan

**Author notes:** Corresponding authors: Wanqiang Qian, Wei Fan.;. Contributed equally (Conghui Liu, Yuwei Ren).

## Abstract

Whole-genome duplication (WGD) has been observed across a wide variety of eukaryotic groups, contributing to evolutionary diversity and environmental adaptability. Mollusks are the second largest group of animals, and are among the organisms that have successfully adapted to the nonmarine realm through aquatic-terrestrial (A-T) transition, and no comprehensive research on WGD has been reported in this group. To explore WGD and the A-T transition in Mollusca, we assembled a chromosome-level reference genome for the giant African snail *Achatina immaculata*, a global invasive species, and compared the genomes of two giant African snails (*A. immaculata* and *Achatina fulica*) to the other available mollusk genomes. The chromosome-level macrosynteny, colinearity blocks, Ks peak and Hox gene clusters collectively suggested the occurrence of a WGD event shared by *A. immaculata* and *A. fulica*. The estimated timing of this WGD event (∼70 MYA) was close to the speciation age of the Sigmurethra-Orthurethra (within Stylommatophora) lineage and the Cretaceous-Tertiary (K-T) mass extinction, indicating that the WGD reported herein may have been a common event shared by all Sigmurethra-Orthurethra species and could have conferred ecological adaptability and genomic plasticity allowing the survival of the K-T extinction. Based on macrosynteny, we deduced an ancestral karyotype containing 8 conserved clusters for the Gastropoda-Bivalvia lineage. To reveal the mechanism of WGD in shaping adaptability to terrestrial ecosystems, we investigated gene families related to the respiration, aestivation and immune defense of giant African snails. Several mucus-related gene families expanded early in the Stylommatophora lineage, functioning in water retention, immune defense and wound healing. The hemocyanins, PCK and FBP families were doubled and retained after WGD, enhancing the capacity for gas exchange and glucose homeostasis in aestivation. After the WGD, zinc metalloproteinase genes were highly tandemly duplicated to protect tissue against ROS damage. This evidence collectively suggests that although the WGD may not have been the direct driver of the A-T transition, it provided an important legacy for the terrestrial adaptation of the giant African snail.

## Introduction

Whole-genome duplication (WGD) is proposed to be a key evolutionary event driving phenotypic complexity, functional novelty, and ecological adaptation ^1, 2^. With the progress of high-throughput sequencing technologies, almost all fundamental lineages of land plants have been characterized as having undergone WGD, while far fewer WGDs among animal lineages have been reported, especially in invertebrate ^3, 4^. Based on chromosome counts and genome size, WGD events have been speculated to have occurred in mollusks, the second largest animal group, which have survived major extinction events and become some of the most successful conquerors of marine, freshwater and terrestrial environments ^5, 6^. Giant African snails, a group of species within Achatinidae possessing unusually large body sizes, have evolved from aquatic ancestors and achieved a pulmonate terrestriality through an aquatic-terrestrial (A-T) transition ^7^. Because of their remarkable ecological adaptability, some members of the giant African snails are considered global invasive species, causing serious damage to agriculture and households ^8^. The success of their colonization of terrestrial ecosystems and adaptation to diverse environments suggests that giant African snails possess advanced genetic plasticity and evolutionary novelties, although the key drivers and underlying mechanisms remain poorly understood.

In contrast to plants, only a few ancient WGDs have been well documented in animals, because polyploidy is usually an evolutionary dead end, resulting from associated meiotic difficulties ^9, 10^. In vertebrates, several ancient WGD events that occurred at the origin of vertebrates and teleost fishes have been proposed to have shaped genetic diversity and adaptive radiation ^11, 12^. In animals other than vertebrates, less conclusive evidence of ancient WGD events has been reported as there is a deficiency of continuous high-quality genome data. Nevertheless, there are several known cases of large-scale genome duplication in groups, such as rotifers, chelicerates and insects. The genome of bdelloid rotifers is tetraploid, and its scaffolds are rearranged during asexual reproduction ^13^. A possible ancestral WGD in chelicerates has been identified in horseshoe crab and was dated to at least 135 million years ago (MYA) through the phylogenetic analysis of Hox genes ^14^, and subsequent surveys in chelicerates revealed that the spider and scorpion lineages underwent another separate WGD before 430 MYA ^15^. Because of the fragmental assembly and uncertain accuracy of traditional analysis methods, the validation of WGD in invertebrates remains controversial. Surprisingly, Li et al. reported 18 ancient WGDs during the evolution of insects in a recent paper, while another macrosyteny analysis in lepidopterans showed that the gene-tree-based and Ks-based detection of WGD adopted by Li are unreliable, and suggested that WGD events should be verified by chromosome-scale genome assembly and macrosynteny analysis ^16, 17^.

Regarding the A-T transition, the conquest of land by organisms that evolved from aquatic ancestors represents one of the most astonishing events in the history of life on Earth ^18^. The dramatic environmental changes from homogeneous water habitat to the heterogenous land environment brought about a significant revolution in species evolution, leading to radiation diversity and life complexity ^19, 20^. This step was achieved in multiple phyla independently via a set of adaptations such as water balance, air breathing, nitrogen excretion, neural-immune system interactions, and certain behaviors ^21^. Within Mollusca, the clade Pulmonata includes several lineages that invaded the terrestrial zone and non-aquatic ecosystems, especially the Stylommatophora ^22^. The giant African snail within Stylommatophora is a representative of the land snails that has also developed fundamental machinery and behaviors such as a pulmonate lung, complex immune system, mucus production, and aestivation, making it highly adapted to the terrestrial environment ^22–24^. Several reports on amphibious species have shown that the expansion and positive selection of genes related to innate immunity, nitrogen excretion, hormonal regulation and pulmonary surfactants were closely connected to the A-T transition ^25^, although the genomic features and evolutionary characteristics of terrestrial invertebrates are still poorly described. Studies of the giant African snail, a promising model for terrestrial mollusks, will facilitate the elucidation of invertebrate A-T transition.

Mollusca is the second largest animal phylum constituting 7% of living animals on Earth ^26^, and comprises numerous species of evolutionary and economic importance ^27, 28^. However, the available genomic resources for Mollusca are still quite insufficient in comparison with those for other large phyla such as Arthropoda and Nematoda ^29, 30^. The genomes of aquatic mollusks such as California sea hare ^31^, Pacific oyster ^32^, pearl oyster ^33, 34^, owl limpet ^35^, California two-spot octopus ^36^, golden mussel ^37^, *Biomphalaria glabrata* ^38^, and golden apple snail ^39^ have been sequenced, whereas few terrestrial species in Mollusca have well-documented genomic information. Recently, the genomic data of the invasive land snail *A. fulica* were released, but without in-depth studies of related biological issues ^40^. To address the genetic and evolutionary characteristics of terrestrial mollusks, we report the genome of another giant African snail with a larger body size and greater invasiveness (Fig. 1a, b and c) ^41^, and provide insights into the molluscan WGD, A-T transition, and invasion mechanisms, through comparative genomic analysis between the two giant African snails and other closely related mollusks.

**Figure 1.**
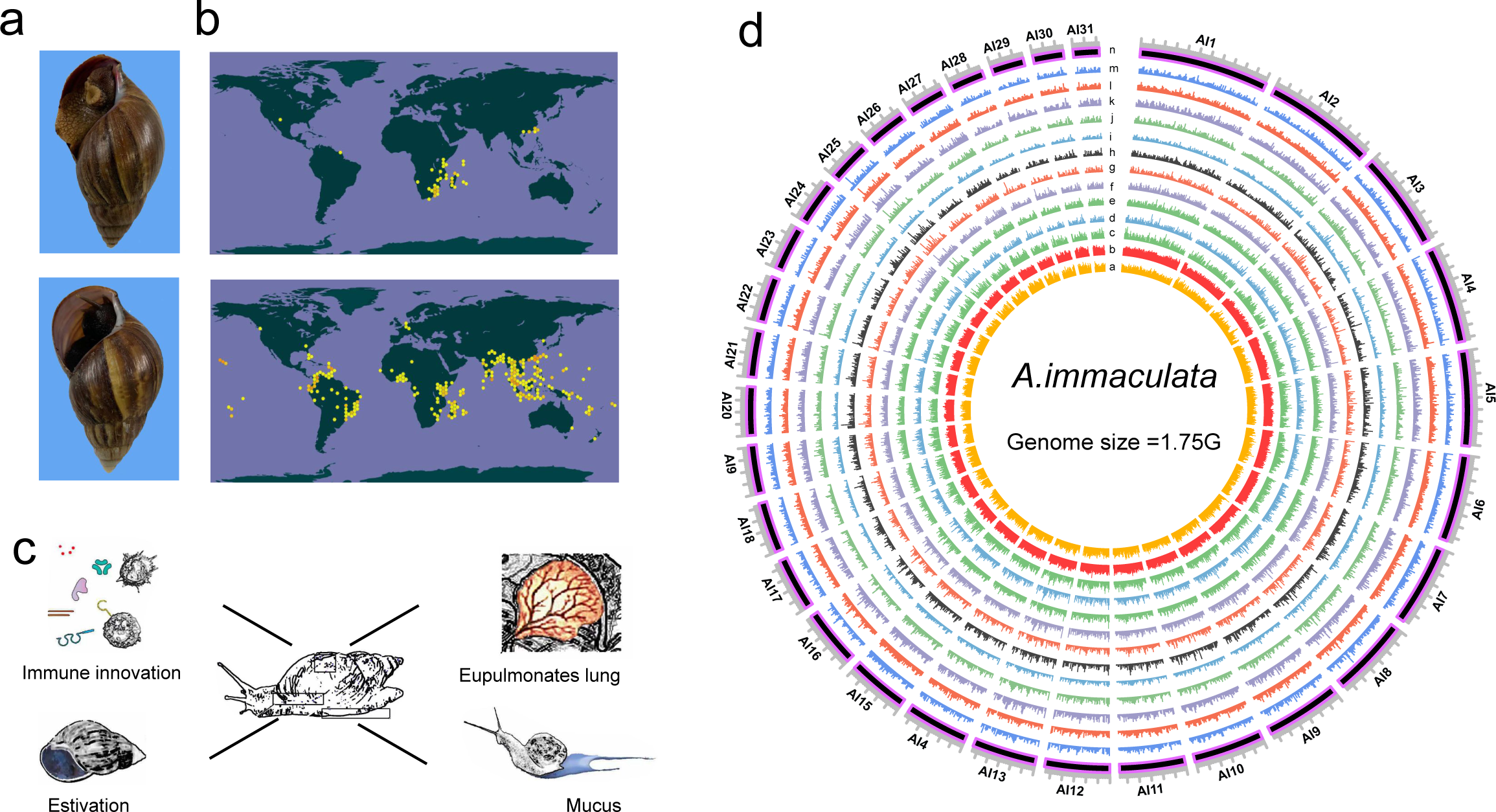
The general characteristics of *A.immaculata*. (**a**) The morphological difference: *A. immaculata* has relatively longer shell with pink or purple columella, while *A. fulica* has shorter shell with white columella. (**b**) Global invasion regions for *A. immaculata* (top) and *A. fulica* (bottom). Currently, *A. fulica* is more widely distributed, but *A. immaculata* has more advantage with a faster invasion speed. **(c)** The sketch map showed the major physiological specialty of *A. immaculata*, including immune innovation, eupulmonates lung, estivation and mucus. **(d)** Genomic features showed by Circos plot. Track n: 31 linkage groups of the genome; Track d-m: expression profile of brain, egg, eye, hemocytes, hepatopancreas, kidney, lungs, muscle, ovary and testis tissues; Track c: distribution of transposon elements; Track b: distribution of gene density; Track a: distribution of GC content.

## Results

### The *A. immaculata* genome provides a better assembly and annotation for giant African snails

We generated 200 Gb (121 ×) of PacBio SMRT raw reads with an average read length of 14 kb, and 145 Gb (82 ×) of Illumina HiSeq paired-end reads using DNA extracted from a single adult of *A. immaculata* (Supplementary Table 1). After quality filtering, 199 Gb (120 ×) of clean PacBio SMRT reads were corrected with Canu ^42^, assembled with wtdbg2 ^43^, and polished with pilon ^44^, resulting in an assembly of 563 raw contigs with a total length of 1,653 Mb and an N50 length of 3.80 Mb. Based on the Hi-C data, 1,648 Mb (99.7%) of final contigs were anchored and arranged into 31 linkage groups, each corresponding to a natural chromosome (Fig. 1d, Supplementary Fig. 1), where the longest was 111.19 Mb and the shortest 34.32 Mb. According to an estimated genome size of 1.75 Gb based on the distribution of the k-mer frequency ^32^ (Supplementary Fig. 2), ∼95% of the *A. immaculata* genome was assembled. To further confirm the accuracy and completeness of the assembly, we mapped the Illumina reads to the assembled reference genome. Significantly, 99.11% of the genome-derived reads could be aligned to the reference genome with a coverage rate of 96.50%, suggesting no obvious bias in sequencing and assembly. The high-quality reference genome provides a good foundation for gene annotation.

Protein-coding genes were predicted in the reference genome by using EVM, integrating *de novo* prediction, transcriptome and homology data (Supplementary Table 2). In total, 28,702 gene models were predicted as the reference gene set, with coding regions spanning ∼39.1 Mb (2.37%) of the genome (Supplementary Table 3 and 4). The distribution of CDS length in *A. immaculata* was similar to that in closely related species (Supplementary Fig. 3 and 4). Overall, 87.5% of the reference genes were supported by transcriptome data, and 96.27% of the eukaryotic core genes from OrthoDB (http://www.orthodb.org/) were identified in the reference gene set by BUSCO. These results were comparable to those obtained from other published molluscan genomes (Supplementary Table 4). In the functional annotation, a total of 26,616 (92.73%) reference genes were annotated according to at least one functional database (Supplementary Fig. 5).

The quality of this assembly is much better than that of other molluscan genomes published thus far (Supplementary Table 4). In particular, the coverage rate and sequence continuity were greatly improved compared with the most recent published *A. fulica* genome. The estimated genome size was 2.12 Gb, and the *A. fulica* genome was assembled into 1.86 Gb. The coverage rate of the *A. immaculata* genome (95%) was 8% higher than that of *A. fulica* (87%). The N50 and N90 lengths of the *A. immaculata* contigs were increased by 5.3 times and 4.9 times, respectively. With better assembly quality, ∼5000 additional gene models were annotated in *A. immaculata*, and the BUSCO rate was improved from 93% (*A. fulica*) to 96% (*A. immaculata*). The high quality of the assembly and annotation of *A. immaculata* provide a better resource for research on giant African snails.

### Signs of adaptive evolution in giant African snails

To gain insights into the evolution of giant African snails (*A. immaculata* and *A. fulica*), a total of 292,034 reference genes from ten molluscan genomes (Fig. 2) were clustered into 17,949 orthologous groups (OGs) containing at least two genes each. Then, a phylogenetic tree was built based on 229 high-confidence single-copy orthologous genes with RAxML ^45^ and the divergence time was estimated using R8S^46^. The results showed that *A. immaculata* diverged from *A. fulica* 21 million years ago (MYA), from *B. glabrata* (Panpulmonata) 174 MYA, from *A. californica* (Euopisthobranchia) 205 MYA, and from *P. canaliculata* (Caenogastropoda) 416 MYA (Fig. 2a). Through CAFE analysis, we identified 2225 expanded OGs in giant African snails (both *A. immaculata* and *A. fulica*). The functions of these OGs are mainly related to signal transduction; the endocrine, immune and nervous systems; longevity regulation and reproduction (Supplementary Fig. 6). Additionally, 836 OGs were found exclusively in the lineage of giant African snails, which mainly functioned in neurohormonal regulation and mucus synthesis and included such as acetylcholine receptor, pannexin, tenascin, adrenocorticotropic hormone receptor, neuropeptide receptor, 5-hydroxytryptamine receptor, mucin, heparan sulfate glucosamine 3-O-sulfotransferase and proteophosphoglycan genes (Fig. 2b, Source Data file). This evidence suggests that the expanded and lineage-specific OGs may play important roles in the adaptation and invasion of giant African snails.

**Figure 2.**
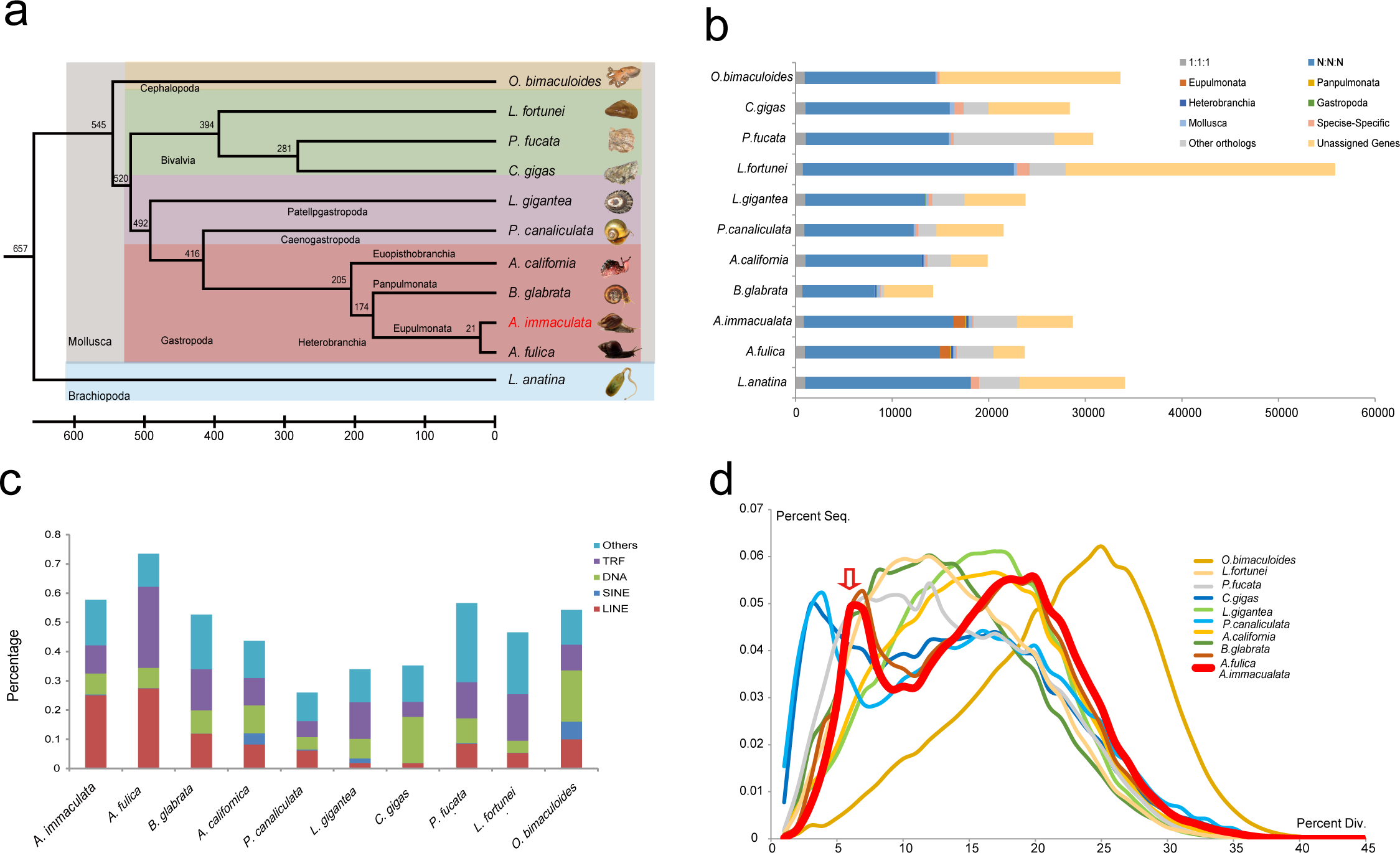
Evolutionary analysis of *A.immaculata*. (**a**) Phylogenetic placement of *A. immaculata* within the dated tree of mollusks. The estimated divergence time is shown at each branching point, and each lineage is shown in different color blocks. *A. immaculata* is highlighted in red. **(b)** Categories of orthologous and paralogous genes in various phyla and species. **(c)** Classification and contents of repetitive sequences in *A. immaculata* and other mollusks. **(d)** Distribution of divergence rate for the class of DNA transposons in mollusks genomes. The divergence rate was calculated by comparing all TE sequences identified in the genome to the corresponding consensus sequence in each TE subfamily. The arrow indicates that both *A. immaculata* and *A. fulica* had a recent explosion of TEs at divergence rate of ∼5%.

This high-quality genome assembly enables a comprehensive analysis of transposable elements (TEs), which play multiple roles in driving genome evolution in eukaryotes ^47^. In total, we identified 954 Mb of repetitive sequences in the assembled *A. immaculata* genome and 1,366 Mb in *A. fulica* (Fig. 2c). Next, we analyzed the divergence rate of each class of TE among the available molluscan genomes. Notably, the TE class of DNA transposons showed a specific peak at a divergence rate of 4∼6% for four invasive species, *A. immaculata, A. fulica, P. canaliculata* and *C. gigas* (Fig. 2d), indicating a recent explosion of DNA transposons. We identified 9,291 genes in region that contained DNA transposons distributed within the specific divergence peak. Based on KEGG annotation, these genes were mainly enriched in signal transduction; the endocrine, immune and nervous systems and reproduction (Supplementary Fig. 7). TEs are powerful facilitators of evolution that generate the “evolutionary potential” to introduce small adaptive changes within a lineage, and the importance of TEs in stress responses and adaptation has been reported in numerous studies ^48, 49^. The recent explosion of DNA TEs in giant African snails could have played important roles in promoting their potential plasticity in stress adaptation.

### Whole genome duplication in the Sigmurethra-Orthurethra branch

Whole-genome duplication (WGD) has rarely been reported in animals, especially in invertebrates, although there is growing suspicions of paleopolyploidy in mollusks based largely on genome sizes and chromosome counts ^6^. With chromosome-level assemblies, we searched for macrosynteny based on homologous gene pairs among four molluscan genomes, from *A. immaculata, A. fulica, P. canaliculata* and *P. fucata*. Our chromosome-level macrosynteny revealed a WGD event shared by two giant African snails, *A. immaculata* and *A. fulica*. The 31 chromosomes of *A. immaculata* could be divided into 14 groups with the preservation of correspondence (Fig. 3a, Supplementary Fig. 8), and the same situation was found in *A. fulica* (Fig. 3b, Supplementary Fig. 8). In *A. immaculata*, 2092 homologous gene pairs with mutual best BLASTP hits were located on the corresponding chromosomes, and this number was 2364 for *A. fulica* (Supplementary Table 5). The comparison of the giant African snails, *P. canaliculata* (n = 14) and *P. fucata* (n = 14) revealed that the karyotype doubled in the lineage leading to giant African snails (Fig. 3c, d and e). The chromosomes of *A. immaculata* and *A. fulica* showed a 1 to 1 corresponding relationship (Fig. 3c, Supplementary Fig. 8), while most chromosomes from *P. canaliculata* (Fig. 3d, Supplementary Fig. 8) and *P. fucata* (Fig. 3d, Supplementary Fig. 8) shared macrosynteny with 2 corresponding chromosomes from giant African snails. The colinearity blocks identified by MCScanX based on BLASTP hits in corresponding chromosome groups, also suggested that WGD events had occurred in two giant African snails (Supplementary Fig. 9 and 10). In the gene age distribution plot of homologous gene pairs, a specific Ks peak shared by *A. immaculata* and *A. fulica* was observed, which was consistent with WGD. The duplication of Hox gene clusters was powerful clue leading to the discovery of ancient WGDs in vertebrates ^50^, so we further compared the giant African snails’ genomes to those with a single Hox cluster and no evidence of WGD. Duplicated Hox gene clusters with specific rearrangements were found in two giant African snail genomes. In conclusion, the macrosynteny, colinearity blocks, Ks peak and Hox gene clusters collectively suggested that a WGD event occurred before the speciation of giant African snails.

**Figure 3.**
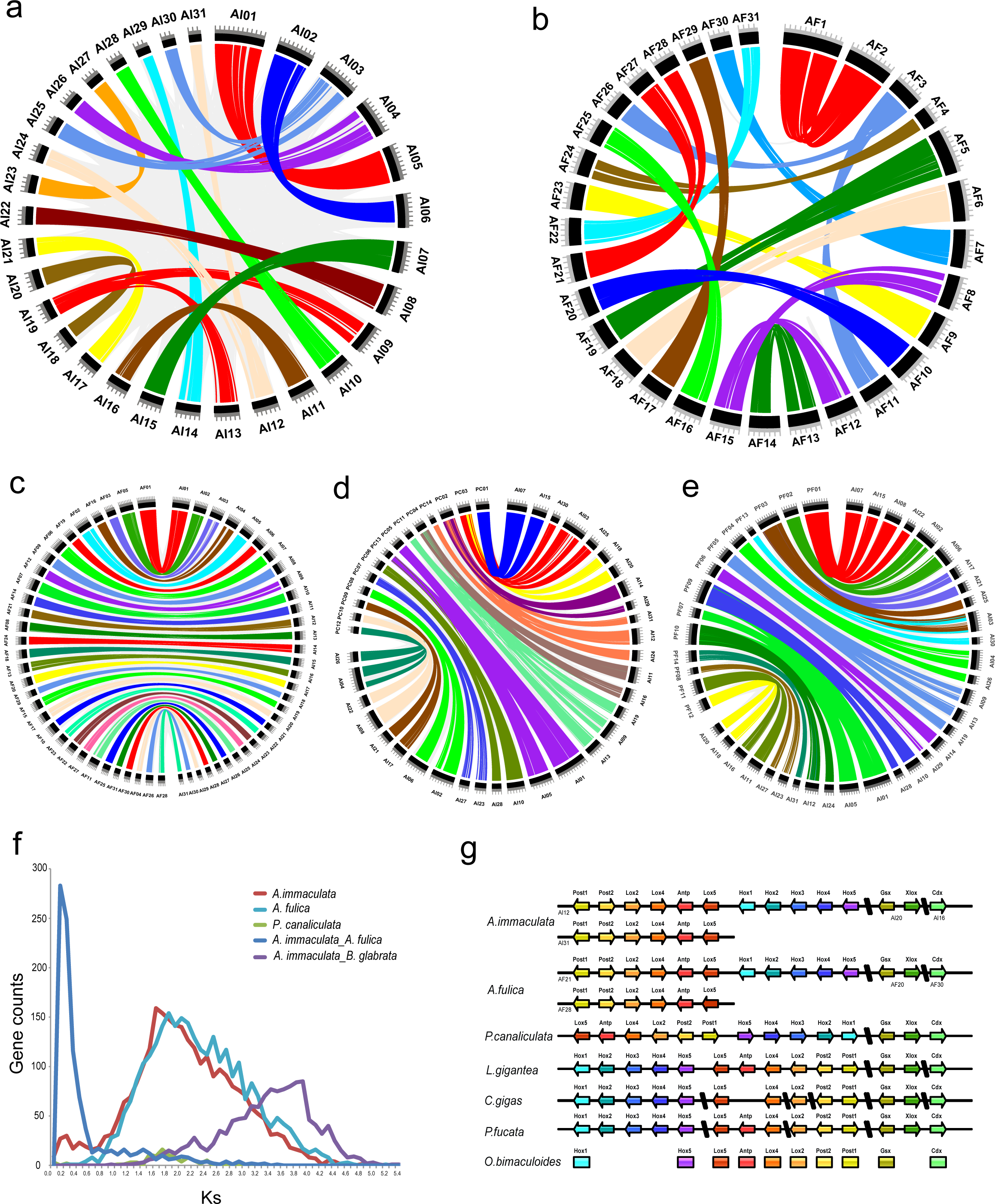
Whole genome duplication shared by *A. immaculata* and *A. fulica.* Circos plots showed the homologous gene pairs in chromosome-level microsynteny among the *A. immaculata* **(a)** and *A. fulica* **(b)** individually, as well as chromosomes between *A. immaculata* and *A. fulica* **(c),** *A. immaculata* and *P. canaliculata* **(d)***, A. immaculata* and *P. fucata* **(e)**. **(f)** Gene age distribution of Ks values calculated from orthologous gene pairs, among *A. immaculata, A. fulica*, *and P. canaliculata* individually, as well as between *A. immaculata and A. fulica*, *A. immaculata and B. glabrata.* **(E)** Comparison of the catalog of Hox genes. *A. immaculata* and *A. fulica* have more Hox genes than the other molloscus, because of the retained genes on the duplicated chromosome. These evidences collectively support a whole genome duplication event shared by *A. immaculata and A. fulica*, which is not found in other mollusks.

**Figure 4.**
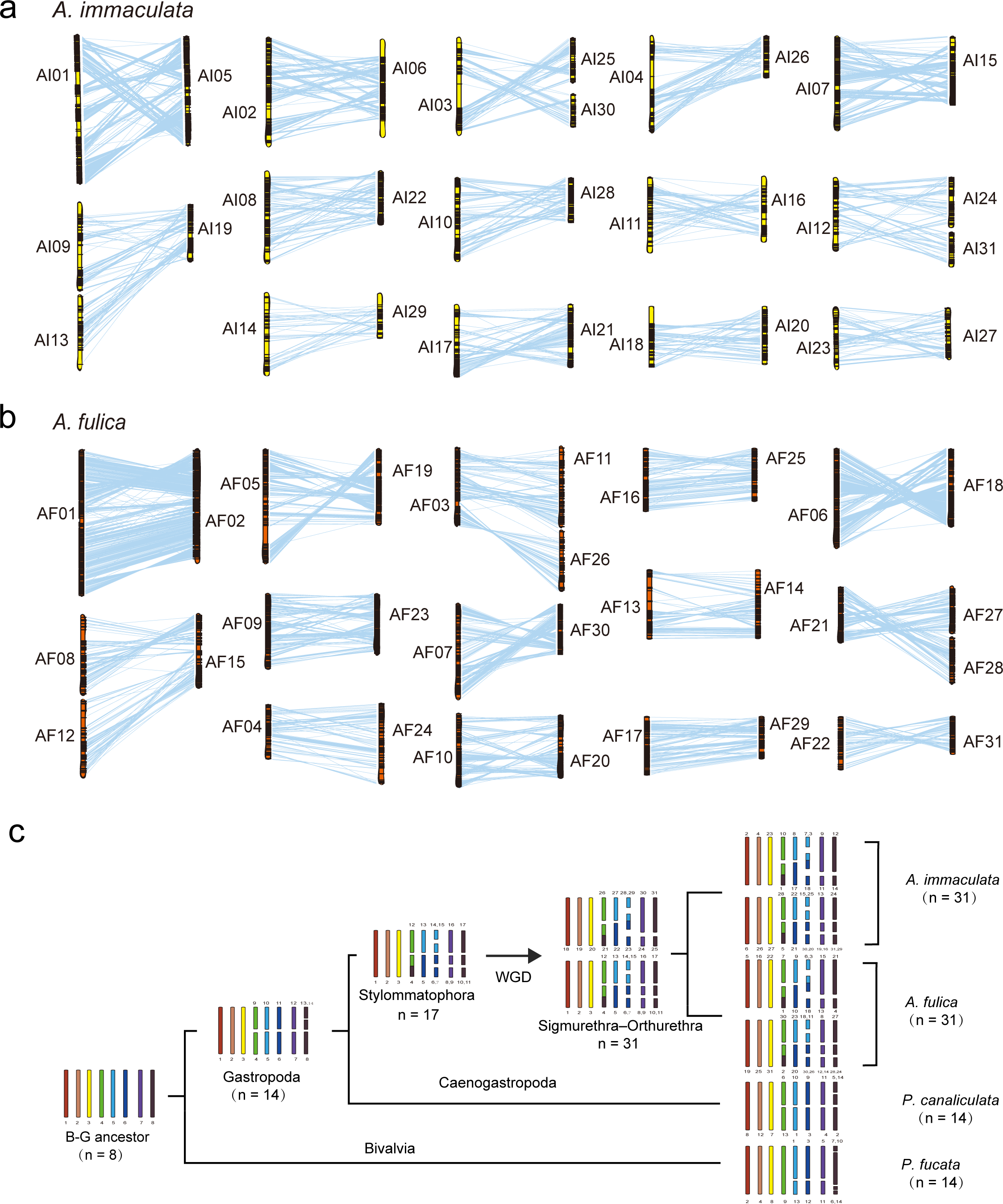
Chromosome relationship and karyotype inferring. Clustering of chromosomes for *A. immaculata* **(a)** and *A. fulica* **(b)**. Each group corresponding to an orthologous chromosome pair derived from WGD, with the links between chromosome pairs representing the mutual best-hit gene pairs. **(c)** Evolution of karyotype in mollusks. The common ancestor of Gastropoda and Bivalvia is estimated to have 8 monoploid chromosomes, while that of Gastropoda has 14. Chromosome breaks, fusions, as well as chromosome number doubling by WGD, formed the current karyotype of living mollusks.

The timing of WGD events has been reported to show a significant correlation with specific geological and global climatic change ^51^. According to the gene age distribution of homologous gene pairs and the divergce time of *B. glabrata* and giant African snails (∼174 MYA), we deduced that the timing of the WGD event was ∼70 MYA. An earlier estimation suggested the occurrence of a WGD event at the Sigmurethra-Orthurethra branch within Stylommatophora by comparison of chromosome numbers among closely related mollusks ^6^. The speciation age of the Sigmurethra-Orthurethra branch was reported to be 65 MYA, which was largely in consistent with our deduced timing, indicating that the WGD reported here was a common event shared by all Sigmurethra and Orthurethra species. The timing of the WGD of the Sigmurethra-Orthurethra branch was also close to Cretaceous-Tertiary (K-T) mass extinction (∼66 MYA), in which global climate change caused the extinction of 60-70% of all plant and animal life, including most mollusks ^52^. In plants, the K-T mass extinction is considered a shared common causal factor in the genome-wide doubling of diverse angiosperm lineages ^53^. It has also been previously suggested that polyploidization in animals is correlated with periods of unstable environments ^1^. The WGD of the Sigmurethra-Orthurethra branch is expected to have provided ecological adaptability and genome plasticity and allowed these taxa to survive the K-T mass extinction.

As the terrestrial area expanded during the K-T mass extinction due to Maastrichtian sea-level regression ^54^, the WGD is expected to have promoted the adaptability of land snails to terrestrial ecosystems and speciation diversity. The functions of WGD-derived homologous gene pairs were significantly enriched in biological regulation, signal transduction, energy generation and the response to stimulus (Supplementary Fig. 11). These functions are closely related to terrestrial living, indicating that the retained WGD-associated genes increased the terrestrial ecological tolerance of giant African snails.

The chromosome-level assembly of *A. immaculata, A. fulica, P. canaliculata* and *P. fucata*, allowed us to infer karyotype evolution within the clade of Gastropoda-Bivalvia based on macrosynteny. Our results indicated a monoploid chromosome number of n=8 for the root of the Gastropoda-Bivalvia lineage. This deduced chromosome number was consistent with that of Patellogastropoda (n=8∼9)^55^, which has been reported to exhibit monophyly in the Gastropoda-Bivalvia lineage and to present preserved ancestral characters ^56^. During subsequent speciation, 6 breaks in the lineages of Gastropoda (n=14) and *P. fucata* (n=14) were observed at different locations. In the Gastropoda clade, Stylommatophora possessed 17 chromosomes, with fusion of chr4 and chr13 of Gastropoda, and showed 4 breaks in chr5, chr6, chr7 and chr8 of Gastropoda, while *P. canaliculata* retained the monoploid chromosome karyotype of Gastropoda (n=14). The WGD in the lineage of the Sigmurethra-Orthurethra branch doubled the chromosome count (n=17+17). After WGD, three fusions (chr6-chr15, chr8-chr16 and chr10-chr17 of Stylommatophora) resulted in the 31 chromosomes of the Sigmurethra-Orthurethra branch (n=17+14) (Fig. 4). In the lineage of *A. immaculata* and *A. fulica*, a chromosome number n = 31 has remained since their speciation in 21 MYA. The demonstration of chromosome duplications, fusions and breaks provides insights into the ancestral karyotype and the mechanism of karyotype evolution in Gastropoda-Bivalvia.

### Expansion of hemocyanins and zinc metalloproteinases improves terrestrial respiratory function

The innovation of respiratory gas exchange is the signature of the aquatic-to-terrestrial transition, which was one of the most conspicuous evolutionary events to have occurred on Earth ^57^. The evolution of the oxygen transportation system allowed land snails to utilize O_2_ from air far more efficiently than aquatic mollusks ^58^. Additionally, an advanced system is needed to eliminate the accompanying oxidative stress to maintain O_2_ homeostasis. However, the underlying mechanisms of O_2_ transport and antioxidation are less well understood.

O_2_ transport in most members of Gastropoda and Cephalopoda is dependent on hemocyanin ^59^. Based on orthologous and phylogenetic analysis, hemocyanin genes were observed in six species of Gastropoda and Cephalopoda, which was consistent with previous reports ^59, 60^. Notably, there are 4 hemocyanin genes in both *A. immaculata* and *A. fulica*, which is twice as many as in any other Gastropoda species (Fig. 5a). Moreover, the four homologous genes of *A. immaculata* and *A. fulica* are located on two chromosomes that derived from WGD, whereas those of other species without WGD were only located in one chromosome or scaffold (Fig. 5b, Supplementary Fig. 12 and 13). Thus, the doubling of the hemocyanin gene number through WGD may have increased the O_2_ transport ability of giant African snails to adapt to land living.

**Figure 5.**
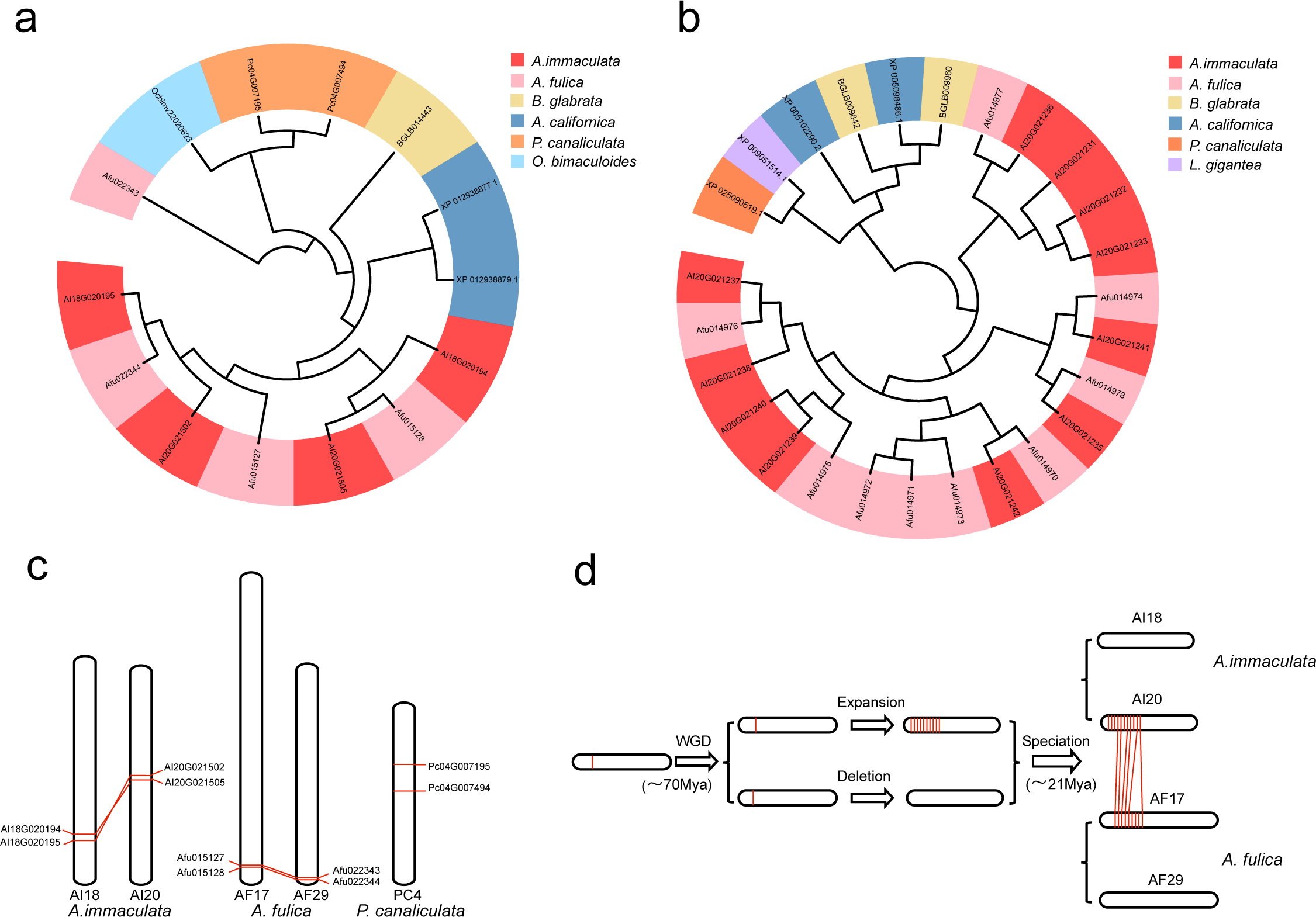
Oxygen transport and anti-oxidization in the respiratory. (**a**) The phylogenetic relationship of hemocyanin genes among mollusks. The protein IDs covered with different colors meant different species. (**b**) The phylogenetic relationship of zinc metalloproteinases among gastropoda, with same style to that of (**a**). (**c**) Comparison of hemocyanin locations in the chromosomes of *A. immaculata*, *A. fulica* and *P. canaliculata*. Hemocyanin genes of *A. immaculata* and *A. fulica* were distributed in two chromosomes, whereas *P. canaliculata* was only in one contig. (**d**) The evolution process of zinc metalloproteinase genes in *A. immaculata* and *A. fulica*. The shared ancient ancestor might have only one zinc metalloproteinase gene located on one chromosome, WGD (∼70 MYA) event doubles the chromosome and gene number, followed by gene expansion on one chromosome and deletion on the corresponding chromosome, resulting in a tandem-cluster of 11 genes and 9 genes in *A. immaculata* and *A. fulica*, respectively.

Reactive oxygen species (ROS) are generated during O_2_ metabolism, and excessive ROS cause oxidative stress and trigger inflammation ^61^. However, antioxidant enzymes protect the host from excess oxidative damage. Superoxide dismutase (SOD), acid phosphatase (AP) and glutathione S transferase (GST) genes were identified in giant African snails, with gene numbers comparable to those of other molluscan species (Supplementary Table 6). In addition to antioxidant enzymes, metalloproteinases have the ability to hydrolyze fibronectin to reduce the injuries caused by ROS ^62, 63^. The number of zinc metalloproteinase genes in both giant African snails was largely expanded, to 11 in *A.immaculata* and 9 in *A. fulica*, compared to only 1 or 2 in other species (Fig. 5c, Supplementary Fig. 14 and 15). Importantly, the zinc metalloproteinase genes of the two species were located in two syntenic clusters on homologous chromosomes, while no homologous genes were found on the corresponding duplicated chromosomes resulting from WGD. The finding that the same location pattern was shared by the two species indicates that these genes were tandemly duplicated after the WGD event but before the specification of *A.immaculata* and *A. fulica*. Furthermore, the expression levels of all zinc metalloproteinases in the hepatopancreas and blood were higher than those in other tissues, which was consistent with the importance of the antioxidative function of these two tissues. Therefore, the expansion of zinc metalloproteinases after WGD might have played important roles in the defense of giant African snails against the damage resulting from oxidative stress and inflammation.

### Glucose homeostasis and ureagenesis benefit survival in aestivation

Aestivation in land animals is a special long-term torpid state that occurs in response to the extreme conditions of summer, such as desiccation, high temperature and starvation ^64^. Terrestrial mollusks originating from aquatic ancestors, developed estivation as a strategy for surviving drastic environmental changes on land. During aestivation, giant African snails seal their shell aperture with epiphragma, and their body remains within the solid shell for several months ^24^. During this long period, these snails are isolated from feeding and excretion, but their blood sugar level and toxic nitrogenous waste are maintained within a normal range. However, how the snails regulate blood glucose homeostasis and eliminate toxins during aestivation is still poorly understood.

Without sugar intake during aestivation, blood glucose homeostasis is mainly achieved via the exploitation of endogenous resources and the reduction of energy expenditure ^65^. Regarding endogenous resources, glycogen is used initially, after which some portion of the carbohydrate skeleton associated with protein metabolism is primarily employed to produce blood glucose through gluconeogenesis ^65^. In this study, we analyzed the expression level changes of two rate-limiting gluconeogenic enzymes phosphoenolpyruvate carboxykinase (PCK, EC:4.1.1.32) and fructose-1,6 - bisphosphatase (FBP, EC:3.1.3.11). Both enzymes are encoded by two homologous genes derived from WGD, and the expression level of one homologous gene is significantly increased in aestivation (negative binomial generalized log-linear model, *p* < 0.01; Supplementary Table 7 and 8), while the other displays relatively constant expression. On the other hand, the consumption of glucose through the tricarboxylic acid (TCA) cycle is minimized in aestivation ^66^. Several important enzymes are involved in the TCA, such as citrate synthase (EC:2.3.3.1), and malate dehydrogenase (EC:1.1.1.37) (Fig. 6a) ^67^. Citrate synthase is encoded by two homologous genes derived from WGD, where one of the homologous genes is significantly downregulated (negative binomial generalized log-linear model, *p* < 0.01; Supplementary Table 7 and 8), while the other gene remains stable. The gene encoding malate dehydrogenase also presented significantly downregulated expression. This evidence indicates that blood glucose homeostasis during aestivation is achieved via both the upregulation of gluconeogenesis and downregulation of the TCA cycle.

**Figure 6.**
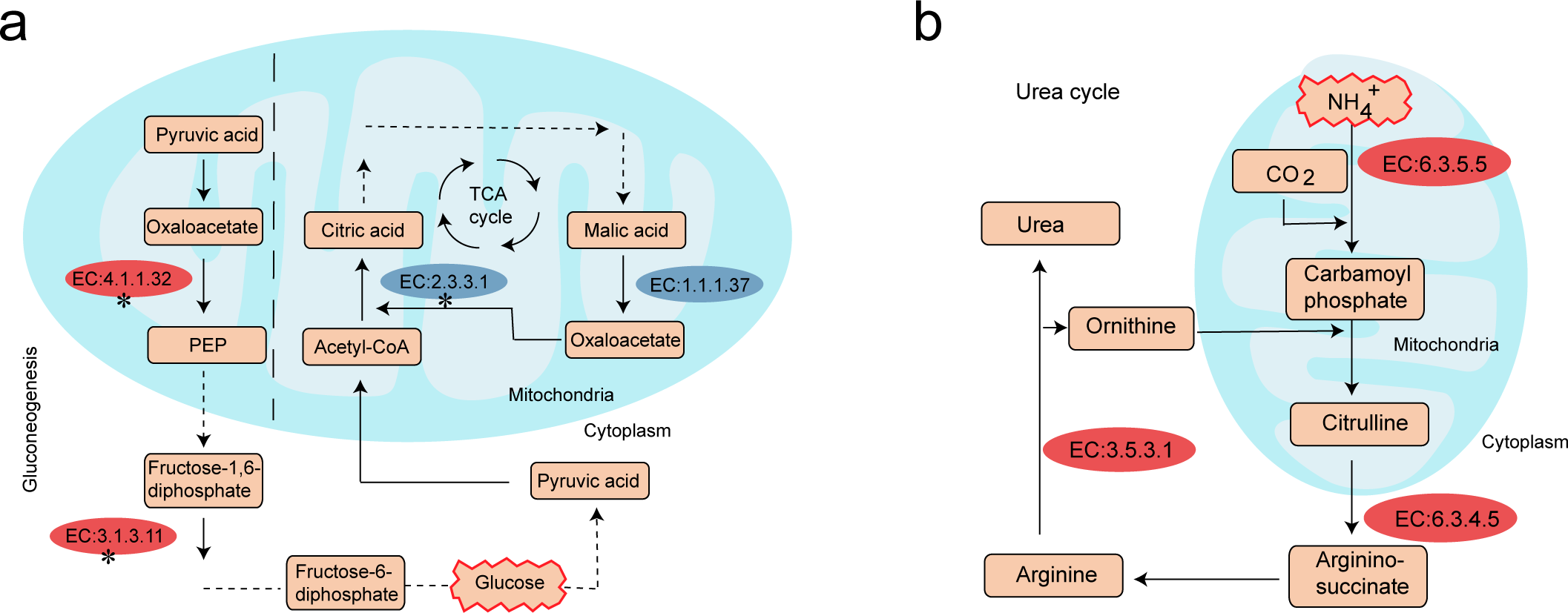
Flowchart of glucose homeostasis and urea-cycle in aestivation of *A. immaculata*. (**a**)The key processes and related gene expression change of gluconeogenesis and TCA cycle in aestivation compared with normal group. The left part represented the major genes participated in gluconeogenesis, while the right part represented the major genes participated in TCA cycle. Two types of arrows were used, double-line arrow meant a number of steps, and single-line arrow meant one step. Two shapes were used, rectangle meant production, and ellipse meant enzymes, with color red and blue representing up-regulated and down-regulated respectively. Ellipse with symbol “*” meant this enzyme had two genes derived from WGD. (**b**) The key processes and related gene expression change of ornithine-urea cycle in aestivation compared with normal groups, the style is same to that of (**a**).

The oxidative deamination of amino acids generates toxic ammonium (NH**_3_**, NH**_4_**^+^), which can cause immunosuppression and neurotoxic effects ^68, 69^. In a normal state, giant African snails mainly convert ammonium into uric acid and excrete it out of their body through the urine, while during aestivation, they convert ammonium into urea and store it in their body ^24, 70^. The accumulation of nontoxic urea during aestivation has been proposed to play a role in the detoxification of nitrogenous substances, the reduction of evaporative water loss, and the reutilization of nitrogen resources ^65^.In this study, we analyzed the expression profiles of three important enzymes involved in the urea cycle, including carbamoyl phosphate synthetase (CPS, EC:6.3.5.5), argininosuccinate synthetase (ASS, EC:6.3.4.5), and arginase (EC:3.5.3.1)^71^. All of the genes encoding these enzymes were significantly upregulated in the aestivation group compared with the normal group (Fig. 6b), providing insight into to the mechanism of the transformation of ammonia into urea during the aestivation of giant African snails.

### Expanded mucus-related gene families reinforce immune defense

Molluscan immune defense has drawn increasing attention because of its economic and evolutionary importance ^5, 72^. As one of the most successful colonizers of terrestrial environments within Mollusca, giant African snails are remarkably adaptive and may possess advanced molecular mechanisms for host-defense against biotic and abiotic stresses ^73^. Although giant African snails lack the canonical vertebrate immunoglobulin, they have developed diverse repertoires of receptors, regulators, and effectors ^23^. To investigate the genes and pathways involved in immune and stress responses, we characterized the immune system on the basis of the genome of *A. immaculata*, including pattern recognition receptors, soluble factors, and mucus-related gene families (Fig. 7a, Supplementary Table 9). The transcriptomes of the hemocytes after different stimuli (lipopolysaccharide/LPS, peptidoglycan/PGN, poly (I:C)/IC and β-glucan/GLU) were also analyzed to address the potential roles of these genes (Supplementary Fig. 16).

**Figure 7.**
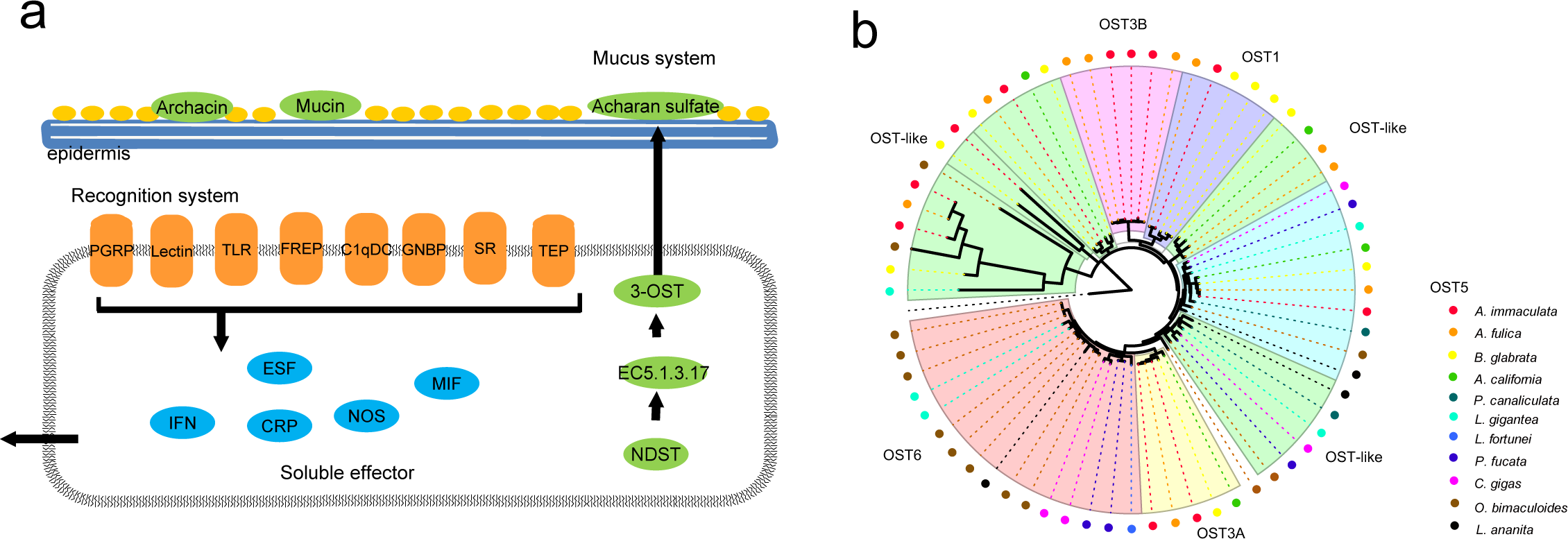
The immune repertoire of *A. immaculata* and the expansion of OST. (**a**) The immunity of *A. immaculata* is presented in mucus, pattern recognition receptors, and soluble effectors. The mucus system covered epidermis, mainly including archacin, mucin and acharan sulfate secreted through the NDST pathway, which were presented with green ellipse. The pattern recognition system on the cytomembrane (orange color) contained PGRP, lectin, TFNR, TLR, FREP, GNBP, SR, TEP, and secreted soluble effector (red ellipse) in the downstream immune cascading. (**b**) The expansion of OST genes. Phylogenetic analyses were performed using MEGA7 through maximum likelihood. The OST genes from different species were indicated in circus filled with different colors.

The mucus of giant African snails is mainly composed of glycosaminoglycans and glycoproteins, and its antimicrobial properties were first reported in the 1980s ^74^. In the *A. immaculata* genome, we identified 11 heparan sulfate glucosamine 3-O-sulfotransferases (OSTs) functioning in glycosaminoglycan synthesis and 99 mucins that encode the major glycoprotein in mucus. OST genes were found to be expanded in Panpulmonata (10 in *A. immaculata*, 11 in *A. fulica*, 10 in *B. glabrata*) and Cephalopoda (13 in *O. bimaculoides*), in which they were two times more abundant than in other mollusks (Supplementary Table 10). The expansion of OSTs in Panpulmonata occurred in subfamilies OST-1 and OST-3B (Fig. 7b). Acharan sulfate, a product of OST, is a primary constituent of mucus, and the expansion of the OSTs was therefore closely related to the abundant surface mucus. The increased expression levels of OSTs observed after LPS stimulus suggest a possible immune role in the response to Gram-negative bacteria. The mucin genes were also found to be expanded on the Stylommatophora branch (99 for *A. immaculate*, 71 for *A. fulica*), showing a number approximately three times greater than in other mollusks, indicating that the mucus of Stylommatophora contains more mucin proteins to defend against microorganism infection (Supplementary Fig. 17, Supplementary Table 10). Increased expression patterns of mucins were mainly detected in the LPS, IC and GLU groups, implying their immune roles against Gram-negative bacteria, viruses and fungi, respectively.

A total of 34 mucins from 19 OGs and 3 OSTs from 2 OGs were found exclusively on the Stylommatophora branch (Source Data file). Their homologous pseudogenes were located within the corresponding duplicated chromosomes, indicating that these genes were generated before WGD, doubled in number through WGD, and transformed into pseudogenes after WGD (Supplementary Table 11 and 12). In addition to their roles in immunity, functions of the mucins and OSTs in wound healing, locomotion and other terrestrial-related processes were observed, indicating that their expansions before WGD may have played important roles in the A-T transition.

## Discussion

WGD provides evolutionary novelties that increase environmental adaptation and species diversity, although this process is regarded as rare in animals because it is hampered by sex determination ^75^. As one of the best-known simultaneous hermaphrodites ^76^, the giant African snail possesses the ability to undergo autofecundation under certain circumstances, thereby overcoming the evolutionary dead end resulting from WGD. Based on two high-quality assemblies from giant African snails, we report a high-credibility WGD on the Sigmurethra-Orthurethra branch (Order: Stylommatophora) with collective evidence including macrosynteny data, colinearity blocks, the gene-age distribution and Hox gene clusters. In particular, chromosome-level macrosynteny was employed for WGD identification, which was highly recommended in a recent paper to avoid possible false-positive results ^17^. To the best of our knowledge, this WGD is the first to be reported in Mollusca and the first to be reported in invertebrates based on chromosome-level macrosynteny. In contrast to the two WGDs found in chelicerates (∼135 and ∼430 MYA), this molluscan WGD was a relatively more recent event with an estimated timing of ∼70 MYA. In contrast to ancient WGDs, the identification of this recent event will help to reveal the adaptive mechanisms and consequences of WGD, especially regarding subgenome divergence, providing more traceable genomic clues ^77^. In most cases, WGD occurs only under remarkably uncommon circumstances and provides a driving force during subsequent evolution. As an example, the WGDs identified in a cluster of angiosperm plants share a common causal factor corresponding with the K-T mass extinction ^78^. The timing of the WGD that occurred in chelicerates at ∼430 MYA and that in Sigmurethra-Orthurethra at∼70 MYA is close to the Ordovician-Silurian (O-S) extinction and the K-T mass extinction, respectively, indicating that the invertebrate WGD was also connected to mass extinction events. Furthermore, it has been reported that WGD is followed by a substantial increase in morphological complexity and species numbers ^2^. Therefore, the species richness and wide ranging adaptations of invertebrates imply that more undiscovered WGDs exist in this clade and might be revealed as the available genomic resources increases.

The A-T transition from water-living to land-living was a milestone in the evolutionary history on Earth ^20^. The Stylommatophora lineage, which originated from a marine ancestor, and breaths through permanently air-filled lungs successfully completed terrestrial adaptation approximately 100∼150 MYA ^79^. With all the associated anatomical and physiological changes required by terrestrial living ^80^, Stylommatophora is considered to provide good study material for the investigation of the A-T transition. The WGD event and sea-level regression that occurred during the K-T boundary are expected to have resulted in greater adaptability to terrestrial ecosystems ^54^. To reveal the mechanism underlying the A-T transition and the influence of WGD, we investigated gene families related to respiration, aestivation and immune defense from the giant African snail. The terrestrial adaptation of Stylommatophora was initiated before WGD (∼70 MYA), and we found that several mucus-related gene families expanded early in the Stylommatophora lineage (100∼150 MYA); these genes included the mucin and OST families, functioning in water retention, immune defense and wound healing. WGD has been proposed to provide functional redundancy and mutational robustness to increase the rates of evolution and adaptation ^81^. The genes encoding hemocyanins, involved in O_2_ transport, and PCK and FBP, involved in gluconeogenesis, were doubled and retained after WGD, enhancing the capacity for gas exchange and glucose homeostasis in aestivation. The extra chromosome copy resulting from genome duplication might limit the risk of genetic variation in one copy of the chromosome. In the post-WGD period, zinc metalloproteinase genes were highly tandemly duplicated, resulting in the protection of tissue against ROS damage arising from respiration. This evidence indicates that although the A-T transition of the giant African snail was not initially driven by WGD, WGD could have facilitated its terrestrial adaptation by providing additional genomic resources, thus increasing the survival rate in the drastic transition from water to land.

Biological invasion has become an increasing serious problem worldwide, as the international flow of people and goods has increased over the years ^82^. With its rich species diversity and strong environmental adaptability, Mollusca is among the phyla with the greatest numbers of invasive species, such as the golden apple snail and giant African snail, which are listed among the top 100 global invasive species, although the invasion mechanism of mollusks is not yet clear. Under genetic bottlenecks, the invasive species can still adapt to new habitats and expand their populations, which is referred to as the ‘genetic paradox of invasive species’ ^83^. Previous reports have shown that TEs are powerful facilitators of rapid adaptation that generate “evolutionary potential” by introducing stress-induced changes in invasive species ^84^. In this study, recent TE explosions were observed in all 4 invasive mollusks but were absent in the other noninvasive mollusks. Genes located near recently emerged TEs were enriched in the function of stress responses, which is consistent with our previous findings in the golden apple snail ^39^, indicating that the recent TE explosion might be a common genetic force contributing to biological invasion. In addition, WGD has been proposed to provide robustness of genetic variation and redundant gene resources for the rapid evolution of novel functions, driving phenotypic complexity and ecological adaptation. Therefore, WGD may be another explanation for the genetic paradox of biological invasion, and species exhibiting WGD may present greater invasiveness.

In summary, we have revealed a WGD occurring on the Sigmurethra-Orthurethra branch within Stylommatophora providing genomic evidence for the A-T transition in Mollusca, and we propose WGD as a potential mechanism contributing to biological invasion. In addition, the obtained genome sequences of giant African snails will enable us to develop more environmentally friendly and efficient control measures using species-specific gene targets, benefiting the protection of agricultural crops and ecological environment as well as the prevention of human disease caused by zoonotic parasites. On the other hand, giant African snails are considered as a high-protein food source in some parts of the world, especially in Africa and Asia. The invasive characteristics of giant African snails, such as their rapid growth, high production rate, and ability to survive harsh conditions, endow them with the potential to be cultivated as an economic species. Thus, the genome sequence of giant African snails provides a powerful platform for the genetic breeding of this species, turning “waste” into wealth.

## Methods

### Samples collection and genome sequencing

Adults of *A. immaculata* were collected from a local farm in Yangjiang, Guangdong province, China, and maintained at 25 ± 2 °C for a week before processing. Genomic DNA was extracted from the foot muscles of a single snail for constructing PCR-free Illumina 350-bp insert libraries and PacBio 20-kb insert library, and sequenced on Illumina HiSeq-X and PacBio SMRT platforms, respectively. The Hi-C library was prepared using the muscle tissue of another single snail by the following methods: Nuclear DNA was cross-linked in situ, extracted, and then digested with a restriction enzyme. The sticky ends of the digested fragments were biotinylated, diluted, and then ligated to each other randomly. Biotinylated DNA fragments were enriched and sheared again for preparing the sequencing library, which was then sequenced on a HiSeq-X platform (Illumina).

### RNA sample preparation and transcriptome sequencing

Ten tissues including brains, eggs (2 days post fertilization), eyes, hemocytes, hepatopancreas, kidneys, lungs, muscles, ovaries and testes from six animals were collected as replicates. Eighty snails were employed for the immune elicitor challenge experiment, and they were equally divided into control group, LPS (lipopolysaccharide) group, PGN (peptidoglycan) group, GLU (β-glucan) group and IC (poly I:C) group. The five groups of snails received injections of 100 μL phosphatic buffer solution (PBS, 0.14 M NaCl, 3 mM KCl, 8 mM NaH_2_PO_4_·12H_2_O, 1.5 mM K_2_HPO_4_, pH7.4), LPS from *Escherichia coli* 0111:B4 (Sigma-Aldrich, 0.5 mg·ml^-1^ in PBS), PGN from *Staphylococcus aureus* (Sigma-Aldrich, 0.8 mg·ml^-1^ in PBS), GLU from *Saccharomyces cerevisiae* (Sigma-Aldrich, 1.0 mg·ml^-1^ in PBS), and poly I:C (Sigma-Aldrich, 1.0 mg·ml^-1^ in PBS), respectively. These treated snails were maintained after injection, and fifteen individuals from each group were randomly sampled at 12 h post-injection. Non-aestivation snails were feeding with enough food and water, whereas aestivation snails were fasting and treated with high temperature, and hemolymph of these two groups were collected around 10 days after the epiphragma formation. Hemolymph samples of all the treated group were collected from five individuals were pooled into one sample. There were three replicates for each sample. In final, total RNAs were extracted from the stored tissues, and then mRNAs were pulled out by beads with poly-T for constructing cDNA libraries (insert 350-bp), and sequenced on an Illumina HiSeq-X sequencer.

### Genome assembly

The Illumina raw reads were filtered by trimming the adapter sequence and low-quality regions (https://github.com/fanagislab/assembly_2ndGeneration/tree/master/clean_illumina), resulting in clean and high-quality reads with an average error rate < 0.001. For the PacBio raw data, the short subreads (< 2 kb) and low-quality (error rate > 0.2) subreads were filtered out, and only one representative subread was retained for each PacBio read. The clean PacBio reads were corrected by Canu 1.8 ^42^ and then assembled by Wtdbg2 ^43^. The PacBio reads were employed to polish the raw contigs by a module within Wtdbg2, after which Illumina reads were aligned to the contigs by BWA-MEM, and single base errors in the contigs were corrected by Pilon 2.10 ^44^ with the parameters “-fix bases, -nonpf, -minqual 20”. Next, Hi-C sequencing data were aligned to the haploid reference contigs by HiC-Pro 2.11.1 ^85^, and then these contigs were clustered, ordered, and orientated into chromosomes with LACH-ESIS ^86^.

### Genome annotation

A de novo repeat library for *A. immaculata* was constructed by RepeatModeler (v1.0.4; http://www.repeatmasker.org/RepeatModeler.html). TEs in the *A. immaculata* genome were identified by RepeatMasker (v4.0.6; http://www.repeatmasker.org/) using both the Repbase library and the de novo library. Tandem repeats in the *A. immaculata* genome were predicted using Tandem Repeats Finder v4.07b ^87^. The divergence rates of TEs were calculated between the identified TE elements in the genome and their consensus sequence at the TE family level.

The gene models in the *A. immaculata* genome were predicted by EVM v1.1.1 ^88^ integrating evidences from ab initio predictions, homology-based searches and RNA-seq alignments. Then, these gene models were annotated by RNA-seq data, UniProt database and InterProScan software ^89^. Finally, the gene models were retained if they had at least one piece of supporting evidence from the UniProt database, InterProScan domain and RNA-seq data. Gene functional annotation was performed by aligning the protein sequences to the NCBI NR, UniProt, COG and KEGG databases with BLASTP v2.3.0+ under an E-value cutoff of 10^-5^ and choosing the best hit. Pathway analysis and functional classification were conducted based on the KEGG database ^90^. InterProScan was used to assign IPR domains and GO terms to the gene models.

### Evolutionary analysis

Orthologous and paralogous groups were assigned from 11 species (*A. immaculata*, *A. fulica, Aplysia californica, Biomphalaria glabrata, Pomacea canaliculata, Lottia giganta, Crassostrea gigas, Pintada fucata, Limnoperna fortune, Octopus bimaculoide* and *Lingula anatina*) by OrthoFinder ^91^ with default parameters. OGs that contained only one gene for each species were selected to construct the phylogenetic tree. The protein sequences of each gene family were independently aligned by muscle v3.8.31 ^92^ and then concatenated into one super-sequence. The phylogenetic tree was constructed by maximum likelihood (ML) using RAxML 8.2.12 ^45^, with the best-fit model (LG+IGF) estimated by ProtTest3 ^93^. The absolute rates of molecular evolution and divergence times were estimated by r8s ^46^. The tree was calibrated with the following time frames to constrain the age of the nodes between the species: minimum = 260 Ma and maximum = 290 MYA for *P. fucata* and *C. gigas* ^94^; minimum = 450 MYA and maximum = 480 MYA for *A. californica* (or *B. glabrata*) and *L. giganta* ^95^. The calibration time (fossil record time) interval (550-610 MYA) of *O. bimaculoides* was adopted from previous results ^96^.

### Transcriptome data analysis

Transcriptome reads were trimmed with the same method for genomic reads (https://github.com/fanagislab/assembly_2ndGeneration/tree/master/clean_illumina), and then mapped to the reference genome of *A. immaculata* using TopHat (v. 2.1.0) with default settings. The expression level of each reference gene in terms of FPKM was computed by cufflinks v2.2.1. A gene was considered to be expressed if the FPKM > 0. Differential gene expression analysis was conducted using cuffdiff v2.2.1.

### WGD verification

The analyses incorporating chromosome-level macrosynteny analysis, colinearity blocks, Ks peak and Hox gene clusters were employed to verify the WGD in giant African snails. Based on 4 chromosome-level molluscan genomes, the macrosynteny was identified using homologous gene sets. Genes from different species would be identified as homologous gene pairs when they had mutual best BLASTP hits with each other. The conserved macrosynteny between species with chromosome-level assemblies was displayed in dot plot. Each dot in the dot plot comparison represents a one-to-one homologous gene pair mentioned above. Based on the dot plot, we inferred the circus plot and dual synteny plot. Then, MCScanX was also used with default parameters to identify the colinearity blocks in *A. immaculata* and *A. fulica* ^97^. Ks distribution of gene pairs in colinearity blocks was calculated by ParaAT ^98^ and KaKs_calculator 2.0 ^99^. The homeobox genes were identified in the giant African snails using BLAST (E < e^−5^) against all homeodomain sequences from the HomeoDB database (http://homeodb.zoo.ox.ac.uk/), and were further confirmed by comparing to the Conserved Domains Database (http://www.ncbi.nlm.nih.gov/cdd).

### Gene family analysis

To identify gene families involved in pathways of respiration, aestivation and immune functions, we performed manual curation for identification of homologous genes by three steps. Initially, we aligned known genes of other close species to the *A. immaculata* genome by BLASTP with best hits (E < e^−5^), and followed by the analysis of paralogous genes performed by OrthoFinder. Then, the obtained genes were used to perform phylogeny analysis by maximum likelihood (ML) method with MEGA7 ^100^, to further validate the accuracy and reveal the phylogenetic relationship of these genes.

## Data availability

Data relating with the findings of this work are available within the paper and the Supplementary Information files. A reporting summary for this Article is available as a Supplementary Information file. Source data are provided as a Source Data file. All the raw sequencing data generated during this study have been deposited at NCBI as a BioProject under accession PRJNA561271. Genomic and transcriptome sequence reads have been deposited in the SRA database with BioSample: SAMN12612888. The Whole Genome Shotgun project of A. immaculata has been deposited at DDBJ/ENA/GenBank under the accession WNKJ00000000. The version described in this paper is version WNKJ00000000. The genome assemblies and annotation files are available at the website ftp://ftp.agis.org.cn/~fanwei/Achatina_immaculata_genome/.

## Code availability

The in-house software clean_adapter [https://github.com/fanagislab/ assembly_2ndGeneration/blob/master/clean_illumina/clean_adapter] and clean_lowqual [https://github.com/fanagislab/assembly_2ndGeneration/blob/master/clean_illumina/ clean_lowqual] are used to filter the adapter and low-quality sequence.

## Acknowledgements

The work was funded by the National key research and development program of China (2016YFC1200600), the Agricultural Science and Technology Innovation Program && The Elite Young Scientists Program of CAAS, Fundamental Research Funds for Central Non-profit Scientific Institution (No.Y2017JC01), the Agricultural Science and Technology Innovation Program Cooperation and Innovation Mission (CAAS-XTCX2016), Fund of Key Laboratory of Shenzhen (ZDSYS20141118170111640), as well as National Natural Science Foundation of China (Grant Nos. 31801804).

## Author contributions

W. F. and W. Q. Q. conceived and led the project; C. H. L. and Y. W. R. prepared DNA and RNA for sequencing; C. H. L. performed genome assembly, annotation, evolution, whole genome duplication, and immune analysis; Y.W.R. performed respiration and aestivation analysis; W. F., W. Q. Q., C. H. L. and Y. W. R. wrote and revised the manuscript.

## Competing interests

The authors declare no competing interests.

