## Supplemental Tables and Figures for "Giant African snail genomes provide insights into molluscan whole-genome duplication and aquatic-terrestrial transition"

**Supplementary tables**

**Supplementary Table 1. Statistics of sequencing data for various types of samples.**

| Sample type | Sequencing Platform | Data Size (reads) | Data Size (base pairs) |
| --- | --- | --- | --- |
| Genome | PacBio RsII | 20,256,202 | 199,427,397,001 |
|  | Hiseq-X  PE150 | 963,975,298 | 144,596,294,700 |
|  | Hi-C | 1,781,034,644 | 267,155,196,600 |
| Transcriptome | Hiseq-X  PE150 | 1,757,315,232 | 261,279,116,187 |

**Supplementary Table 2. Summary of transcriptome data and mapping rate.**

| Sample | Total read numbers | Total bases (bp) | Mapping rate (%) |
| --- | --- | --- | --- |
| Brn1 | 42,430,264 | 6,330,009,686 | 81.33 |
| Brn2 | 43,216,366 | 6,452,076,419 | 85.72 |
| Brn3 | 13,638,012 | 2,036,454,181 | 86.42 |
| Egg1 | 42,876,592 | 6,376,000,696 | 97.4 |
| Egg2 | 43,062,758 | 6,401,626,331 | 98.16 |
| Egg3 | 25,338,780 | 3,764,019,925 | 98.14 |
| Eye1 | 43,189,620 | 6,439,912,770 | 90.92 |
| Eye2 | 42,927,742 | 6,400,742,576 | 92.08 |
| Eye3 | 16,327,274 | 2,434,290,064 | 91.17 |
| HEM1 | 29,814,930 | 4,452,980,509 | 51.41 |
| HEM2 | 30,169,492 | 4,508,819,300 | 57.42 |
| HEM3 | 30,390,022 | 4,542,869,893 | 58.33 |
| Hep1 | 41,244,874 | 6,161,720,972 | 67.01 |
| Hep2 | 41,731,948 | 6,237,662,425 | 70.91 |
| Hep3 | 41,790,332 | 6,246,453,266 | 71.44 |
| Kny1 | 41,324,580 | 6,168,313,466 | 65.71 |
| Kny2 | 41,821,362 | 6,245,613,979 | 70.06 |
| Kny3 | 27,037,746 | 4,038,953,495 | 70.36 |
| Lun1 | 40,811,534 | 6,088,404,752 | 71.04 |
| Lun2 | 41,457,016 | 6,188,798,533 | 73.4 |
| Lun3 | 15,757,276 | 2,352,795,293 | 73.1 |
| Mus1 | 43,471,062 | 6,483,885,708 | 91.49 |
| Mus2 | 43,204,096 | 6,443,497,184 | 93.03 |
| Mus3 | 29,847,302 | 4,452,284,138 | 92.59 |
| OV1 | 42,786,254 | 6,361,689,720 | 57.48 |
| OV2 | 43,002,730 | 5,576,118,432 | 97.84 |
| OV3 | 20,780,530 | 3,086,303,537 | 97.67 |
| TE1 | 42,963,804 | 6,394,035,947 | 95.91 |
| TE2 | 42,958,616 | 6,388,289,472 | 97.11 |
| TE3 | 17,990,724 | 2,675,998,857 | 96.82 |
| GLU1 | 43,490,916 | 6,489,285,978 | 84.37 |
| GLU2 | 29,056,448 | 4,335,550,913 | 84.92 |
| GLU3 | 39,448,466 | 5,887,788,055 | 89.52 |
| IC1 | 37,720,524 | 5,626,199,245 | 81.86 |
| IC2 | 42,111,740 | 6,282,762,889 | 82.52 |
| IC3 | 41,941,186 | 6,259,377,577 | 79.49 |
| LPS1 | 42,814,384 | 6,382,338,302 | 90.14 |
| LPS2 | 42,504,474 | 6,335,984,114 | 90.48 |
| LPS3 | 27,519,510 | 4,103,211,600 | 91.07 |
| PGN1 | 39,943,564 | 5,963,512,033 | 76.76 |
| PGN2 | 39,076,530 | 5,835,419,959 | 87.45 |
| PGN3 | 40,332,238 | 6,020,551,142 | 77.84 |
| Aes1 | 38,728,274 | 5,780,019,642 | 83.03 |
| Aes2 | 36,534,992 | 5,454,799,399 | 82.17 |
| Aes3 | 39,330,688 | 5,871,112,081 | 83.69 |
| P1 | 33,899,128 | 5,057,242,188 | 78.03 |
| P2 | 34,490,214 | 5,145,478,473 | 75.42 |
| P3 | 45,008,318 | 6,717,861,071 | 76.72 |

Note：Ten tissues (Brn, brain; Egg, egg; Eye, eye; Hem, hemocytes; Hep, hepatopancreas; Kny, kidney; Lun, lungs; Mus, muscle; Ov, Ovary; Te, testis), 6 hemocyte samples after treatment (GLU, glucan; IC, poly I:C; LPS, lipopolysaccharide; PGN, peptidoglycan; Aes, Aestivation; P, PBS). The transcriptome data was mapped to the genome of *A. immaculata* by Tophat. The mapping rates vary across tissues, from the highest 98.16% in Egg to lowest 51.41% in HEM. The low mapping rate in some tissues may be caused by the low quality of extracted RNA or some unknown contaminations. However, even if the HEM samples with the lowest mapping rate still has over 50% mapped reads, which is enough for the downstream analysis.

**Supplementary Table 3. Statistics of gene predictions from different software in *A. immaculata.***

| Statistics | AUGUSTUS | GeneWise  *P.canaliculata* | GeneWise  *L. gigantea* | RNA-seq | EVM |
| --- | --- | --- | --- | --- | --- |
| Gene counts | 23,152 | 9,512 | 5,268 | 58,324 | 28,702 |
| Avg. CDS length | 1,132 | 846 | 619 | 1,350 | 1,363 |
| Avg. exon number | 4.24 | 4.10 | 2.80 | 3.13 | 6.22 |
| Number of exon | 98,085 | 39,022 | 14,748 | 182,625 | 178,656 |
| Avg. intron length | 1,894 | 3,534 | 3,685 | 239 | 3,052 |
| Avg. exon length | 267 | 206 | 221 | 431 | 219 |
| Single exon gene rate | 0.24 | 0.52 | 0.66 | 0.59 | 0.16 |

**Supplementary Table 4. Summary of assembly and annotation of mollusc genomes.**

| Genome feature | Assembled sequences (bp) | Contig N50 size (bp) | Contig N90 size (bp) | Scaffold N50 size (bp) | Scaffold N90 size (bp) | GC content (%) | No. of gene models | Avg. CDS Length (bp) | BUSCO (%) |
| --- | --- | --- | --- | --- | --- | --- | --- | --- | --- |
| *A. immaculata* | 1,653,153,977 | 3,802,429 | 698,996 | 56,367,627 | 34,315,726 | 38.9 | 28,702 | 1,363 | 96.3 |
| *A.*  *fulica* | 1,855,892,613 | 721,038 | 141,756 | 59,589,303 | 44,109,545 | 38.8 | 23,726 | 1,560 | 93 |
| *B.*  *glabrata* | 916,377,450 | 18,978 | 5,132 | 48,059 | 817 | 36 | 14,224 | 1,066 | 72.8 |
| *A.*  *california* | 927,310,431 | 9,817 | 1,626 | 917,541 | 207,390 | 40.3 | 19,909 | 1,568 | 98.7 |
| *P. canaliculata* | 440,071,717 | 1,072,857 | 303,904 | 31,531,291 | 23,662,357 | 40.3 | 21,533 | 1,497 | 98.9 |
| *L.*  *gigantea* | 359,505,668 | 94,165 | 10,180 | 1,870,055 | 74,480 | 33.3 | 23,824 | 1,136 | 98.4 |
| *C.*  *gigas* | 557,735,934 | 37,218 | 11,109 | 401,685 | 68,181 | 33.4 | 28,402 | 1,472 | 99.4 |
| *O. bimaculoides* | 23,381,887,882 | 5,982 | 1,606 | 475,182 | 79,088 | 36 | 33,638 | 1,535 | 98.7 |

**Supplementary Table 5. Summary of homologous gene pairs of *A. immaculata* and *A. fulica* identified through chromosome-level macrosyteny.**

|  | WGD | TD | Others | TOTAL |
| --- | --- | --- | --- | --- |
| AI_AI | 2,092 | 1,278 | 1,760 | 5,130 |
| AF_AF | 2,364 | 1,319 | 1,367 | 5,050 |
| AI_AF | 11,150 | 0 | 2,495 | 13,645 |
| AI_PF | 3,835 | 0 | 2,889 | 6,724 |
| AI_PC | 6,719 | 0 | 1,812 | 8,531 |

Note: The numbers of homologous gene pairs with mutual best BLASTP hits were indicated as “TOTAL”. The numbers of homologous gene pairs located in WGD corresponding chromosomes were indicated as “WGD”. The numbers of homologous gene pairs located in identical chromosomes were deemed as tandem duplicated and indicated as “TD”. The numbers of homologous gene pairs dispersed in random chromosome were indicated as “others”.

**Supplementary Table 6. Antioxidant enzymes identified in *A. immaculata.***

| Gene name | Gene ID |
| --- | --- |
| Acid phosphatase | AI04G005587, AI04G005598, AI05G007208, AI17G018937, AI08G010632, AI02G002713, AI02G002712, AI02G002034, AI22G022871, AI18G019671, AI06G007580, AI01G000038, AI01G001709 |
| Gst | AI18G019897, AI03G003436, AI03G003435, AI03G003434, AI03G003433, AI03G003432, AI02G002841, AI04G005422, AI25G024679, AI05G006000, AI03G004038, AI10G012526, AI21G021916, AI21G022392, AI12G014075, AI12G014076, AI12G014077, AI06G007318, AI06G007317, AI06G007316, AI06G007315, AI06G007314, AI19G020516, AI04G005286, AI04G005124, AI04G005122 |
| Sod | AI10G012140, AI28G026752, AI10G012536, AI28G026844, AI28G026939, AI10G012752 |

**Supplementary Table 7. Genes related with glucose homeostasis and ureagenesis in aestivation.**

| Function | KEGG_EC_ID | Gene name | *A.immaculata*_ID |
| --- | --- | --- | --- |
| Gluconeogenesis | EC:4.1.1.32 | Phoenolpyruvate carboxykinase | AI04G004789 |
|  |  |  | AI26G025497 |
|  | EC:3.1.3.11 | Fructose-1,6-bisphosphatase | AI05G006172 |
|  |  |  | AI01G001647 |
| TCA cycle | EC:2.3.3.1 | Citrate synthase | AI05G006192 |
|  |  |  | AI01G001591 |
|  | EC:1.1.1.37 | Malate_dehydrogenase | AI02G001873 |
| Ureagenesis | EC:6.3.5.5 | Carbamoyl phosphate synthetase | AI30G027817 |
|  | EC:6.3.4.5 | Argininosuccinate synthetase | AI13G015440 |
|  | EC:3.5.3.1 | Arginase | AI27G026284 |
|  |  |  | AI03G003968 |

**Supplementary Table 8. Expression of glucose homeostasis and ureagenesis related genes in aestivation.**

| *A.immaculata*_ID | P1_  FPKM | P2_  FPKM | P3_  FPKM | AES1_  FPKM | AES2_  FPKM | AES3_  FPKM | FC(fold change) | *p*-value |
| --- | --- | --- | --- | --- | --- | --- | --- | --- |
| AI04G004789 | 76.292 | 57.292 | 95.291 | 63.570 | 46.915 | 80.225 | 1.376 | 0.0799 |
| AI26G025497 | 18.014 | 12.580 | 23.449 | 66.170 | 49.153 | 83.188 |  | 5.00E-05 |
| AI05G006172 | 3.776 | 1.198 | 6.353 | 1.386 | 0.000 | 2.891 | 1.422 | 0.0171 |
| AI01G001647 | 37.669 | 26.491 | 48.847 | 57.570 | 41.490 | 73.649 |  | 0.00025 |
| AI05G006192 | 9.187 | 5.490 | 12.884 | 9.275 | 5.497 | 13.052 | 0.782 | 0.9522 |
| AI01G001591 | 81.001 | 60.064 | 101.937 | 61.252 | 44.591 | 77.913 |  | 0.009 |
| AI02G001873 | 889.960 | 664.524 | 1115.400 | 655.724 | 489.786 | 821.663 | 0.737 | 0.00295 |
| AI30G027817 | 4.121 | 2.724 | 5.519 | 57.239 | 43.053 | 71.424 | 13.888 | 5.00E-05 |
| AI13G015440 | 54.381 | 39.745 | 69.016 | 373.264 | 279.531 | 466.997 | 6.864 | 5.00E-05 |
| AI27G026284 | 7.323 | 3.992 | 10.653 | 55.374 | 40.176 | 70.573 | 25.309 | 5.00E-05 |
| AI03G003968 | 0.526 | 0.000 | 1.928 | 150.653 | 112.053 | 189.254 |  | 0.0002 |

Note: P means normal group, and AES means aestivation group. The FC (fold change) is the average expression of aestivation group divided by that of the normal group. The *p*-value was calculated by the statistic method of negative binomial generalized log-linear model invoked in Cuffdiff.

**Supplementary Table 9. Genes identified in the immune system of *A. immaculata***

| Constituent part | Gene name | Gene ID |
| --- | --- | --- |
| PRR | Lectin | AI24G024028, AI03G003784, AI24G024024, AI08G009964, AI05G007129, AI22G022999, AI22G022995, AI22G022994, AI27G025936, AI17G018934, AI25G024835, AI13G015830, AI12G014393, AI02G003144, AI02G003146, AI29G027418, AI05G006959, AI09G011335, AI06G007969, AI06G007957, AI06G007956, AI09G011373, AI09G011368, AI29G027671, AI05G007031, AI27G026014 |
|  | TLR | AI13G015576, AI13G015366, AI19G021064, AI10G012926, AI10G012924, AI10G012978, AI10G012982, AI10G012983, AI05G007269, AI05G006349, AI07G009390, AI25G025136, AI00G028663, AI19G020384, AI12G014282, AI12G014283, AI12G014284, AI11G013807, AI03G004046, AI10G012528, AI12G014724, AI18G019571, AI10G012626, AI13G015578, AI10G012400, AI28G027105, AI28G027102, AI13G015464, AI13G015465, AI01G001198, AI10G012551, AI21G022441, AI10G012535, AI28G026740, AI12G014979, AI12G014981, AI13G015580 |
|  | C1qDC | AI07G009473, AI13G015831, AI03G003497, AI06G008485 |
|  | FREP | AI25G025256, AI25G024846, AI25G024843, AI25G024832, AI25G024803, AI25G025226, AI25G025225, AI25G024852, AI26G025623, AI25G025251, AI25G025212, AI25G025215, AI25G024849 |
|  | GNBP | AI04G005990, AI04G005888, AI04G005889, AI04G005890, AI26G025261 |
|  | PGRP | AI21G022502, AI21G022501 |
|  | SR | AI16G017902, AI17G019168, AI27G025912, AI22G022856, AI20G021735, AI27G026175, AI20G021851, AI26G025279 |
|  | TEP | AI18G020159, AI18G019744, AI20G021824, AI20G021822, AI18G019749 |
| Soluble factors | CRP | AI08G010572 |
|  | ESF | AI04G005653, AI07G009195, AI08G010657, AI08G010658, AI31G028412, AI04G005960, AI04G005967, AI15G017093, AI17G019317, AI04G005871, AI15G017279, AI01G000399 |
|  | MIF | AI16G018480, AI15G017749, AI16G018294 |
|  | IFN | AI10G012143, AI22G022939 |
|  | iNOS | AI16G018477, AI16G018478, AI11G013274, AI11G013271, AI11G013270, AI11G013269, AI11G013268, AI11G013264 |
| Mucus | achacin | AI09G011426, AI19G021088, AI19G021117 |
|  | EC5.1.3.17 | AI14G016903, AI14G016179, AI29G027364 |
|  | NDST | AI11G013293, AI11G013291, AI11G013289, AI07G008675 |
|  | OST | AI15G017088, AI07G009459, AI08G010303, AI15G017089, AI07G009457, AI04G005828, AI08G009946, AI02G002900, AI27G026450, AI27G026451, AI27G026452 |
|  | mucin | AI03G003689, AI04G005628, AI13G015937, AI19G021076, AI03G004486, AI06G008572, AI10G012380, AI07G009648, AI03G004133, AI11G013193, AI01G001106, AI01G001105, AI02G002939, AI05G007217, AI22G022686, AI11G013362, AI11G013361, AI26G025467, AI19G020677, AI13G015499, AI05G006363, AI15G017518, AI12G014231, AI12G014311, AI09G011258, AI09G011257, AI09G011256, AI09G011255, AI09G011254, AI09G011253, AI09G011252, AI09G011239, AI09G011238, AI06G007559, AI30G027775, AI30G027797, AI30G027838, AI23G023643, AI19G020838, AI10G012572, AI31G028643, AI14G016570, AI01G001056, AI22G022889, AI10G012111, AI05G006895, AI16G018357, AI06G007696, AI13G015355, AI30G028145, AI12G014078, AI03G003596, AI31G028343, AI02G002369, AI30G027723, AI30G027722, AI29G027526, AI29G027525, AI20G021149, AI05G006526, AI05G007064, AI20G021251, AI02G002463, AI12G014583, AI27G026026, AI08G010103, AI03G003714, AI29G027558, AI29G027618, AI16G017868, AI12G014148, AI17G019157, AI29G027298, AI27G025958, AI04G005401, AI05G006848, AI25G024839, AI08G010714, AI06G008023, AI04G004615, AI02G002020, AI30G027767, AI13G015642, AI03G004264, AI06G007661, AI13G015673, AI11G013694, AI01G000017, AI27G026329, AI06G008219, AI23G023469, AI09G011130, AI04G005243, AI02G002773, AI22G023113, AI10G012743, AI09G010974, AI09G011953, AI27G026024 |

**Supplementary Table 10. Gene numbers of mucin and OST (heparan sulfate glucosamine 3-O-sulfotransferase) in mollusc genome.**

| Molluscs | Mucin gene number | OST gene number |
| --- | --- | --- |
| *A. immaculata* | 99 | 11 |
| *A. fulica* | 71 | 10 |
| *B. glabrata* | 22 | 10 |
| *A. californi* | 36 | 4 |
| *P. canaliculata* | 18 | 2 |
| *L. gigantea* | 23 | 5 |
| *C. gigas* | 18 | 4 |
| *P. fucata* | 17 | 4 |
| *L. fortunei* | 42 | 1 |
| *O. bimaculoides* | 24 | 13 |
| *L. ananita* | 20 | 3 |

**Supplementary Table 11. Position and orientation of pseudo OST (heparan sulfate glucosamine 3-O-sulfotransferase).**

| Stylommatophora specific OST ID | Position of pseudo OST | Orientation of pseudo OST |
| --- | --- | --- |
| AI27G026450 | - |  |
| AI27G026451 | Chr23:14699193-14699339 | - |
| AI27G026452 | Chr23:21641174-21641794 | + |

**Supplementary Table 12. Position and orientation of pseudo Mucin.**

| Stylommatophora specific Mucin ID | Position of pseudo Mucin | Orientation of pseudo Mucin |
| --- | --- | --- |
| AI01G000017 | Chr5:45919964-45920467 | - |
| AI02G002020 | Chr6:23841440-23842057 | + |
| AI02G002773 | Chr6:64438410-64438772 | + |
| AI03G004264 | Chr25:25661507-37267185 | - |
| AI03G003714 | Chr30:1175465-1202063 | - |
| AI04G004615 | Chr26:16769064-21429980 | - |
| AI04G005401 | Chr26:33157704-33157811 | + |
| AI05G006848 | Chr1:79136906-79137598 | - |
| AI06G007661 | Chr2:13871669-13872268 | + |
| AI06G008023 | Chr2:9720031-9720249 | - |
| AI06G008219 | Chr2:40732094-40733002 | + |
| AI08G010714 | Chr22:30658494-37394255 | - |
| AI09G011130 | Chr13:16474052-16474507 | - |
| AI09G011953 | Chr13:20098057-20098578 | - |
| AI10G012743 | Chr28:24277790-28244851 | + |
| AI13G015642 | Chr9:10649002-10649694 | - |
| AI13G015673 | Chr9:20336338-25442244 | - |
| AI16G017868 | Chr11:42666469-42667362 | - |
| AI17G019157 | Chr21:17929481-17930636 | - |
| AI22G023113 | Chr8:4322201-4322479 | + |
| AI23G023469 | Chr27:7841909-14848266 | - |
| AI25G024839 | Chr3:25903974-25905317 | - |
| AI27G025958 | Chr23:18905991-18906452 | - |
| AI27G026024 | Chr23:21259851-21261311 | - |
| AI27G026329 | Chr23:14392574-14393488 | - |
| AI29G027298 | Chr14:31902065-48582725 | + |
| AI29G027618 | Chr14:25196430-25196981 | - |
| AI30G027767 | Chr3:8029430-8029864 | + |
| AI04G005243 | - |  |
| AI08G010103 | - |  |
| AI09G010974 | - |  |
| AI11G013694 | - |  |
| AI12G014148 | - |  |
| AI29G027558 | - |  |

**Supplementary figures**

**Supplementary Fig. 1. A genome-wide contact matrix of *A*. *immaculata* from Hi-C data.** The color value indicates the base 2 logarithm of the number of valid reads (log2[valid reads]). Chromosome id are labeled from 1 to 31.

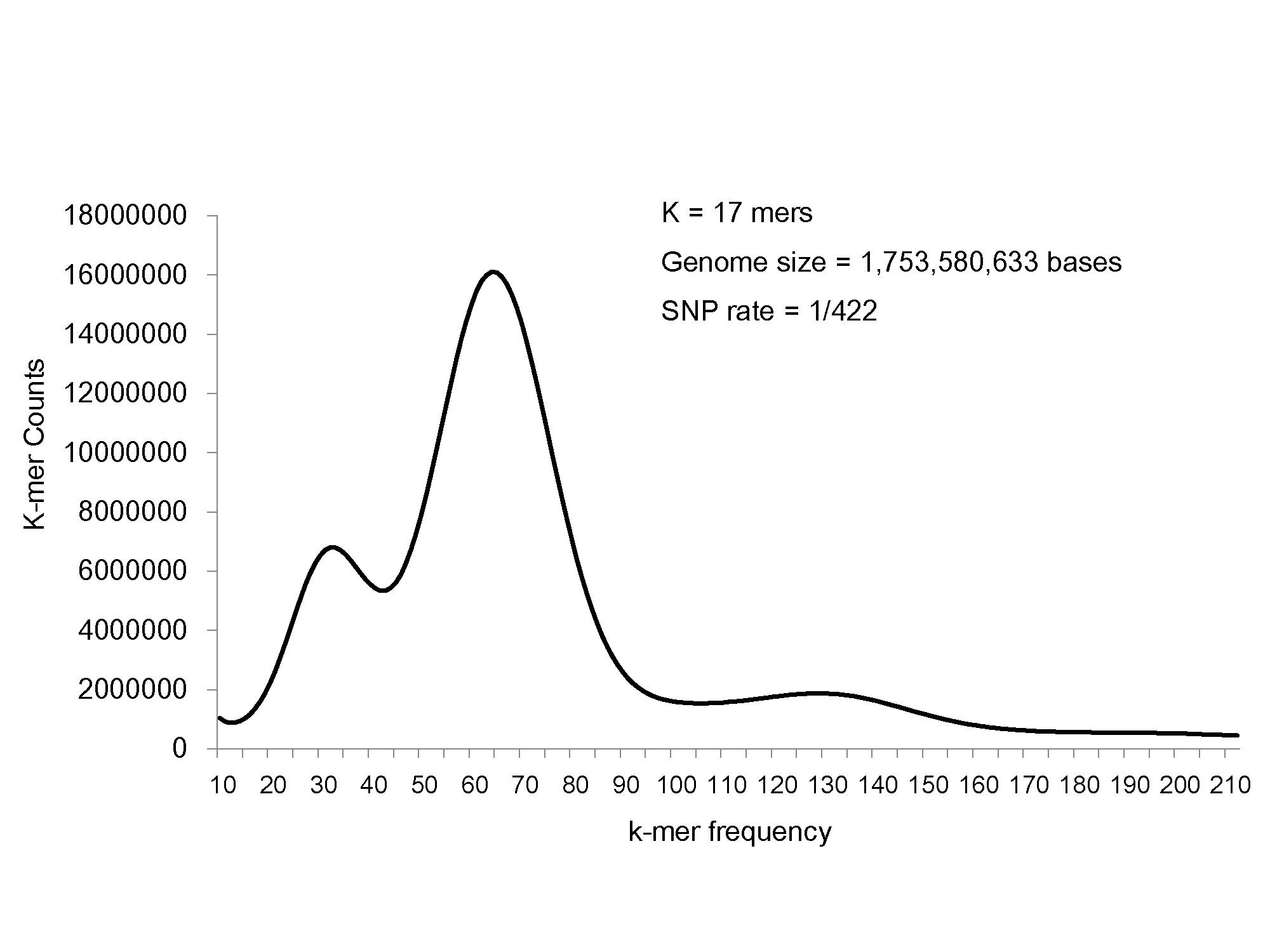
 **Supplementary Fig. 2. A genome survey of *A*. *immaculata* based on Illumina data.** SNP rate was calculated by FindErrors, from AllPaths-LG (Gnerre S, MacCallum I, et al. 2011)

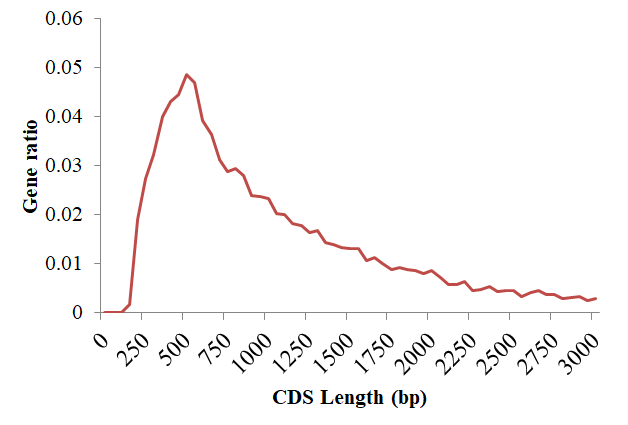

**Supplementary Fig. 3. Distribution of CDS length in *A. immaculata.***

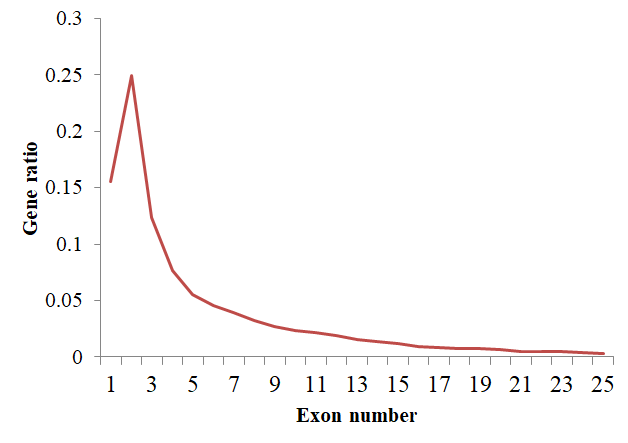

**Supplementary Fig. 4. Distribution of exon number in *A. immaculata*.**

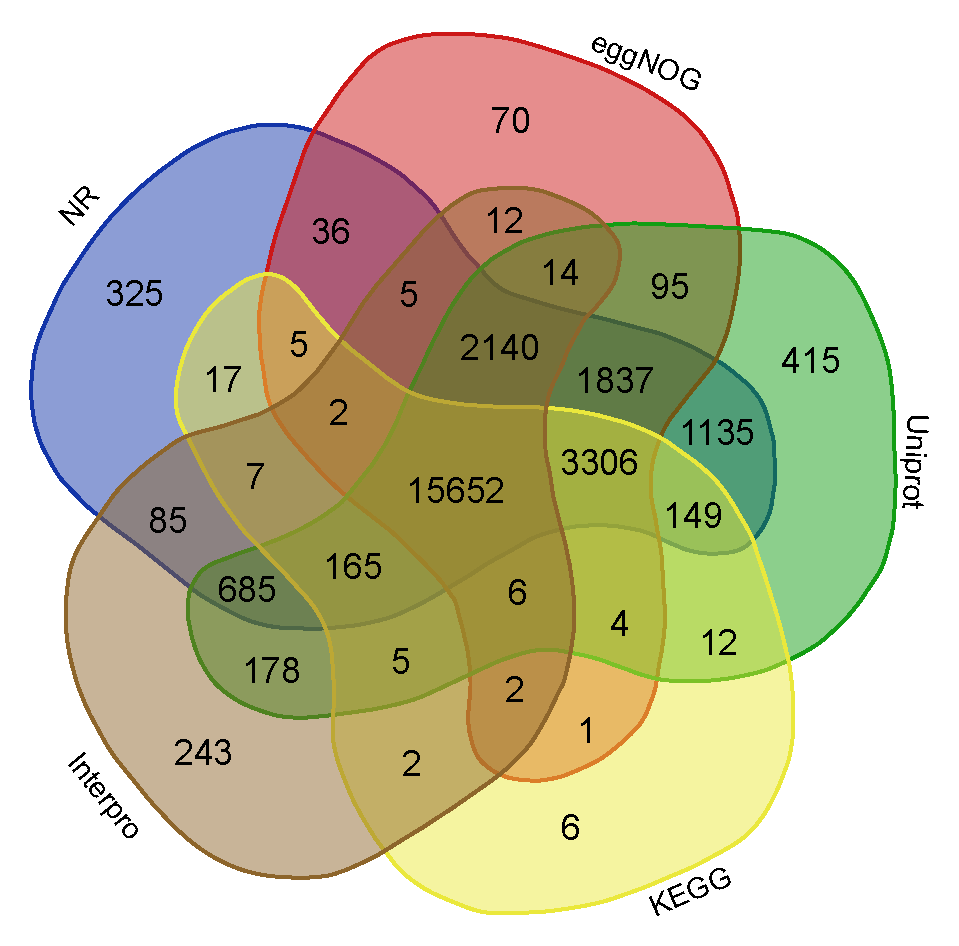

**Supplementary Fig. 5. Reference genes annotated by the eggNOG, KEGG, NR, InterPro and UniProt databases.**

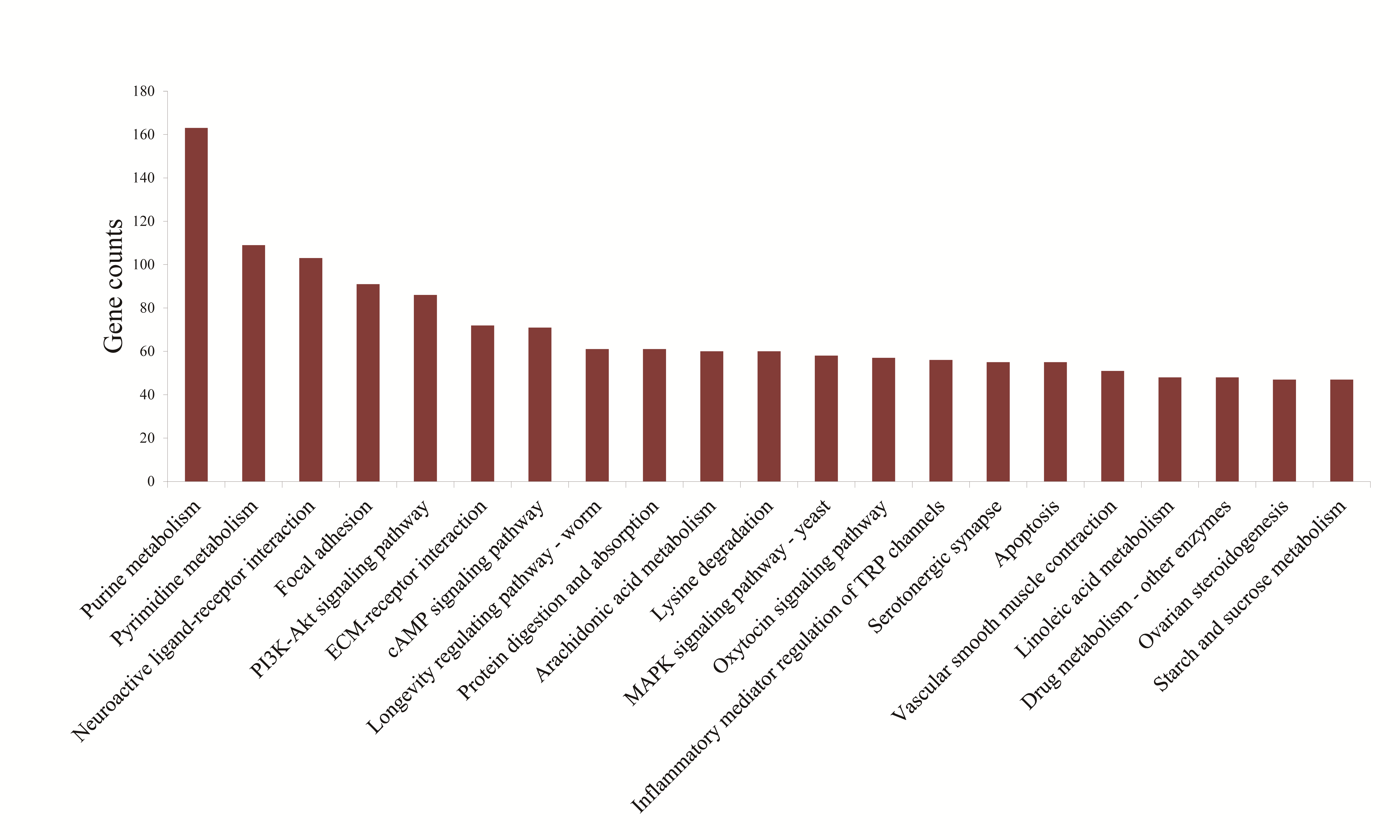

**Supplementary Fig. 6. Function of expanded OGs in *A. immaculata* annotated by KEGG.**

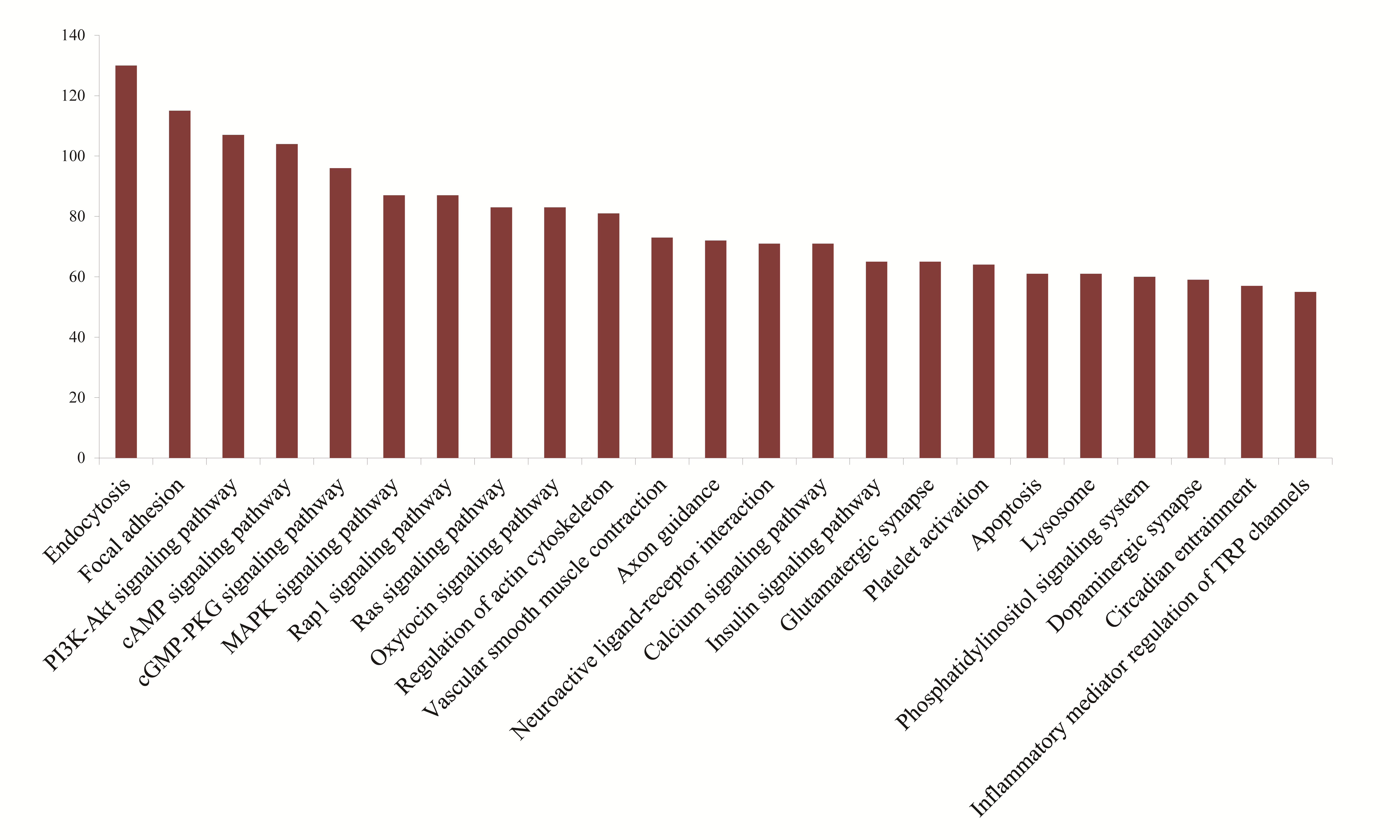

**Supplementary Fig. 7. KEGG function of genes containing DNA transposons distributed within the specific divergence peak.**

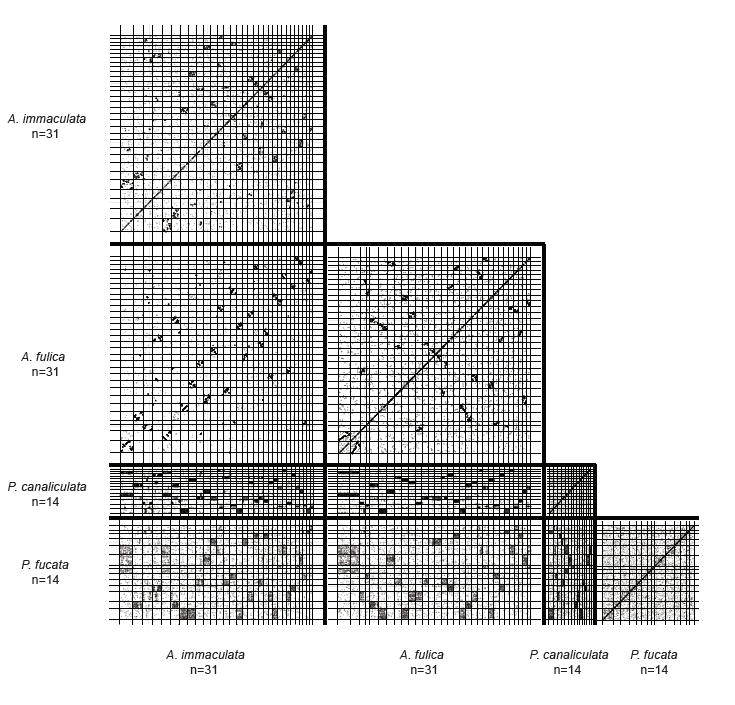

**Supplementary Fig. 8. Chromosome-based macrosynteny is shown in the form of dot plots with comparisons among *A. immaculata, A. fulica, P. canaliculata* and *P. fucata.***

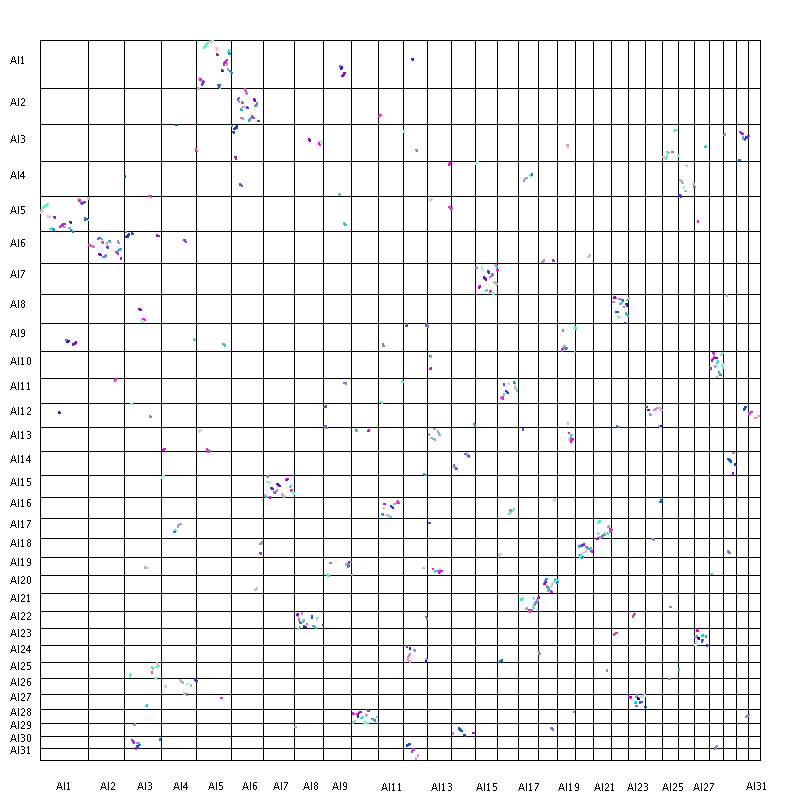

**Supplementary Fig. 9. The colinearity blocks of *A. immaculata* identified by MCScanX based on BLASTP hits.**

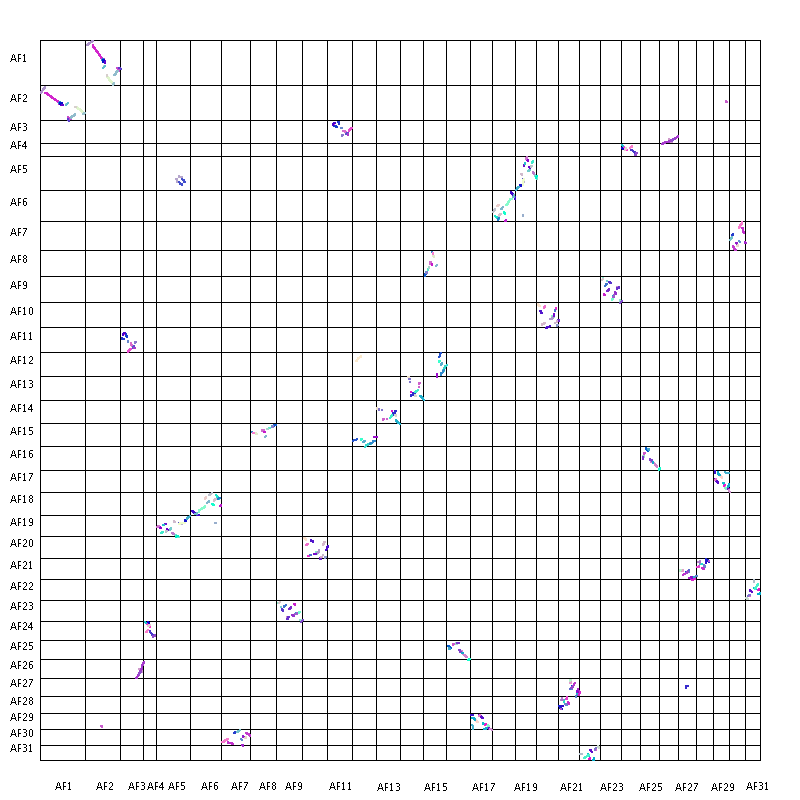

**Supplementary Fig. 10. The colinearity blocks of *A. fulica* identified by MCScanX based on BLASTP hits.**

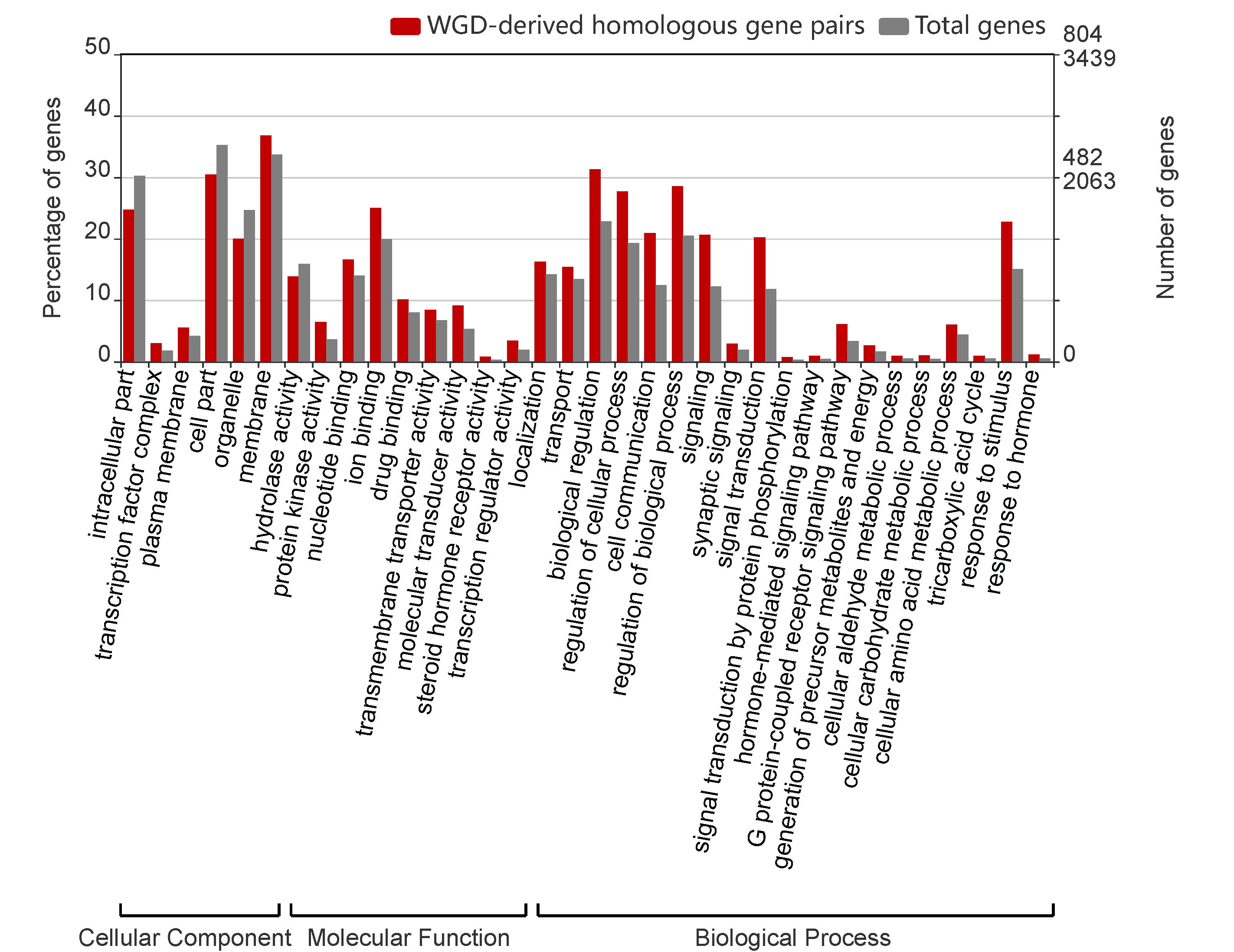

**Supplementary Fig. 11. The enriched GO functions of WGD-derived homologous gene pairs.**

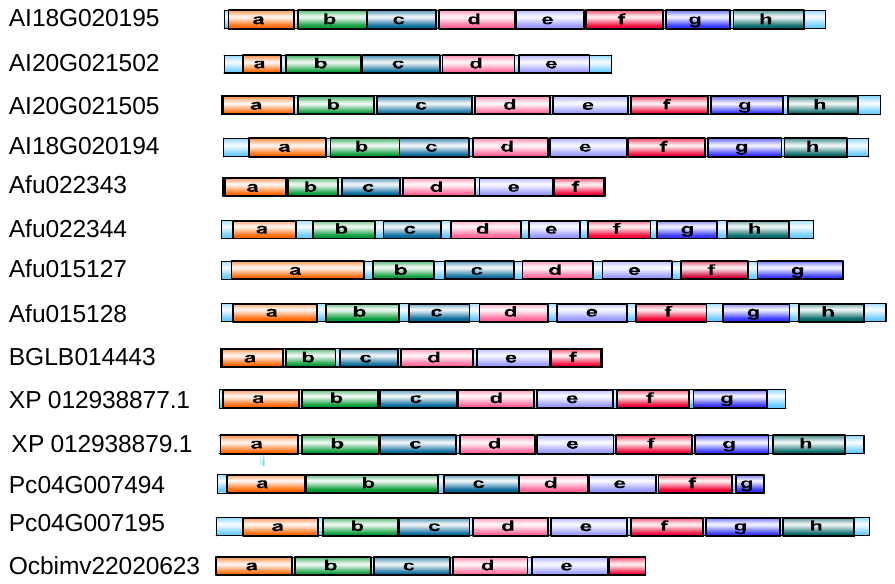

**Supplementary Fig. 12. Function units of hemocyanins in five gastropoda species and *O. bimaculoides.*** Different rectangles meant different function units, the light blue rectangle meant whole length of the protein, the letter of a-h represented eight different function units, gastropoda species and cephalopoda with eight and six function units were annotated as intact protein, respectively. The genes were from six species, A. immaculata, AI18G020195, AI18G020194, AI20G021502, AI20G021505; A. fulica, Afu015127, Afu015128, Afu022343, Afu022344; Biomphalaria glabrata BGLB014443; Aplysia californica, XP_012938877.1 and XP_012938879.1; Pomacea canaliculata, Pc04G007195, Pc04G007494; Octopus bimaculoides, Ocbimv22020623m.p.

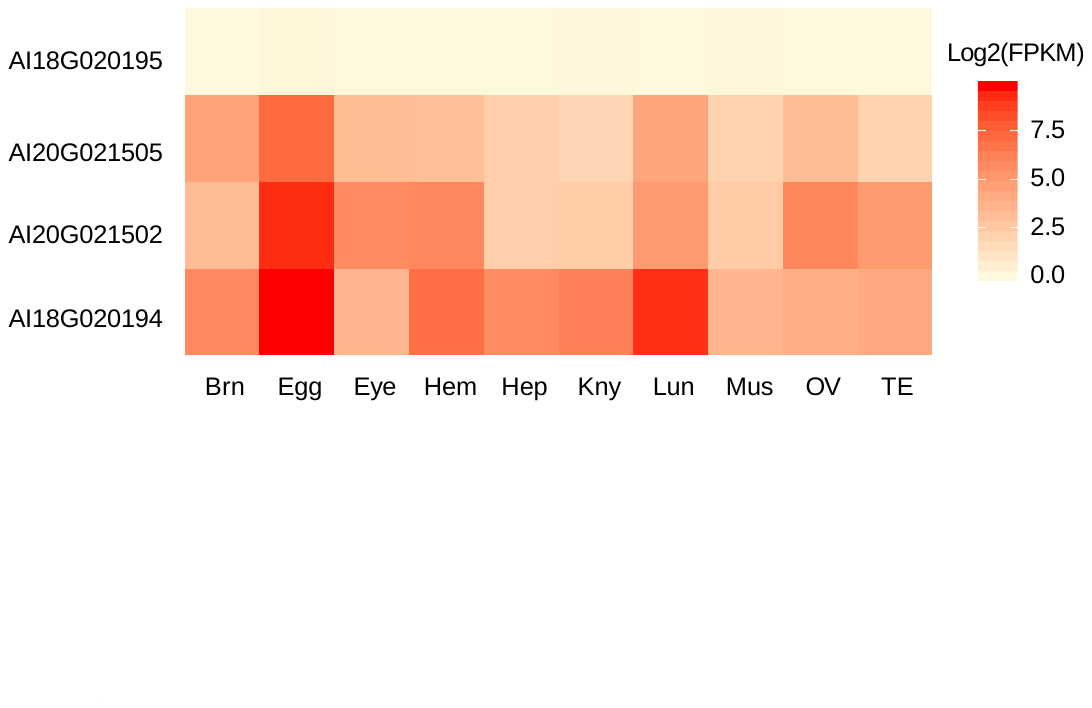

**Supplementary Fig. 13. Expression of hemocyanins in different tissues for *A. immaculata*.** Average expression of 4 hemocyanin genes in ten tissues (Brn, brain; Egg, egg; Eye, eye; Hem, hemocytes; Hep, hepatopancreas; Kny, kidney; Lun, lung; Mus, muscle; Ov, Ovary and albumen gland; Te, testis). Three hemocyanin genes highly expressed in all tissues, whereas AI18G020195 expressed extremely low in any tissue.

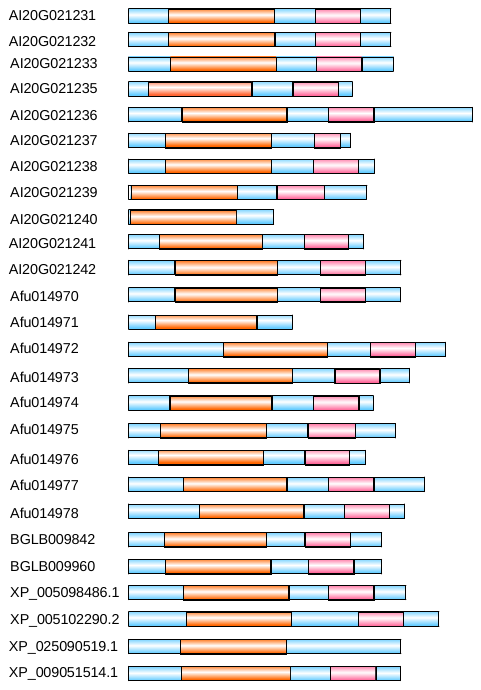

**Supplementary Fig. 14. Functional domains of zinc metalloproteinases in gastropoda species.** Different rectangles represent different domains, the light blue rectangle represents whole length of the protein, orange rectangle represents the metallopeptidase domain, and pink rectangle represents ShKT domain. The genes were from six species, *A. immaculata*, AI20G021236, AI20G021231, AI20G021232, AI20G021233, AI20G021241, AI20G021235, AI20G021242, AI20G021239, AI20G021240, AI20G021238, AI20G021237; *A. fulica,* Afu014977, Afu014974, Afu014978, Afu014970, Afu014973, Afu014971, Afu014972, Afu014975, Afu014976; *B. glabrata*, BGLB009842, BGLB009960; *A. californica*, XP 005102290.2, XP 005098486.1; *P. canaliculata*, XP 025090519.1; *L. gigantea*, XP 009051514.1.

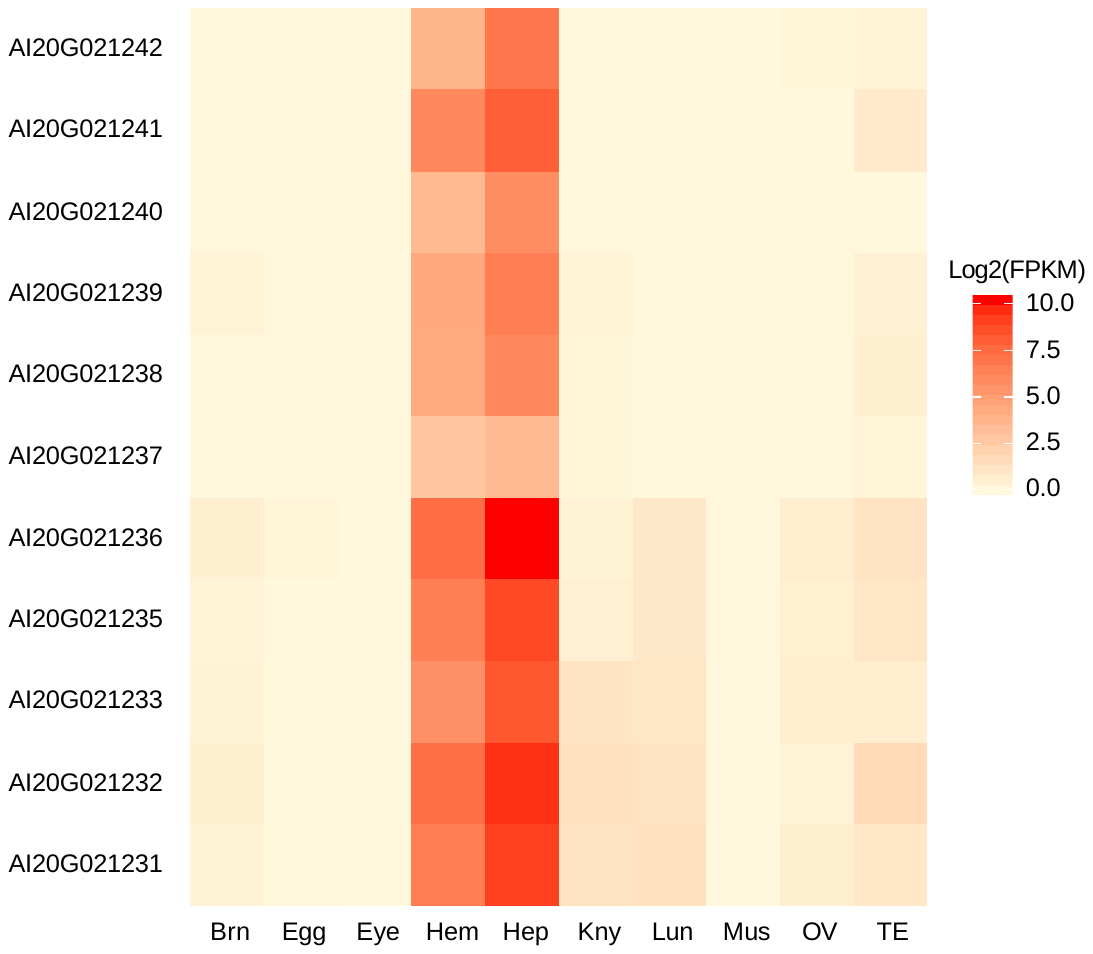

**Supplementary Fig. 15. Expression of zinc metalloproteinases in different tissues for *A. immaculata*.** Average of expression of 11 zinc metalloproteinases genes in ten tissues (Brn, brain; Egg, egg; Eye, eye; Hem, hemocytes; Hep, hepatopancreas; Kny, kidney; Lun, lung; Mus, muscle; Ov, Ovary and albumen gland; Te, testis). All the zinc metalloproteinases genes showed higher expression in Hep and Hem than other tissues. Previous reports showed hepatopancreas and hemocytes are two vital antioxidant tissues, indicating the important roles of these genes in antioxidation.

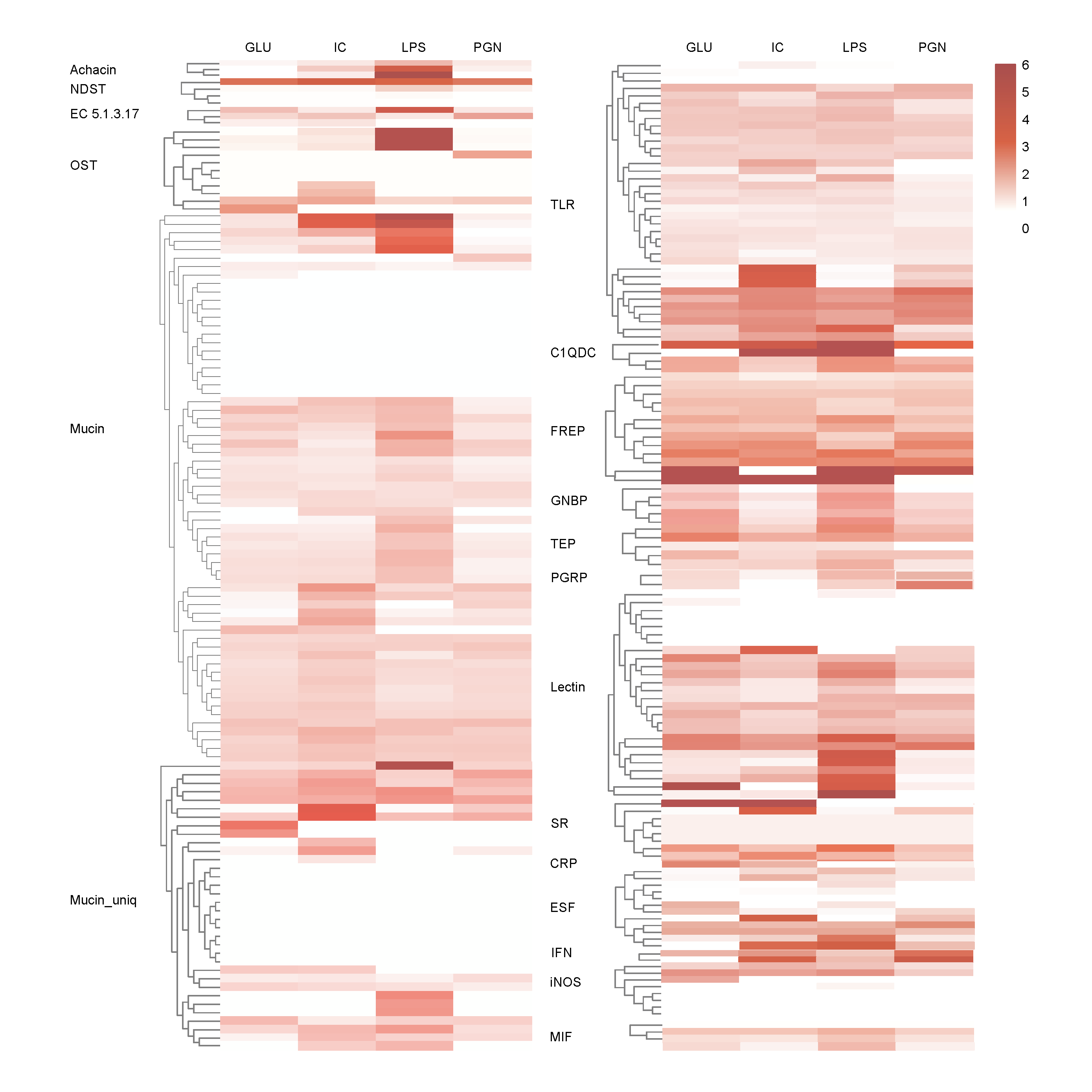

**Supplementary Fig. 16. The expression patterns of the immune repertoire of *A. immaculata* after GLU, IC, LPS and PGN stimuli.** The color value indicates the induced fold change (log2(FPKM-stimulus/FPKM-control)) under the immune stimulus of GLU，IC，LPS and PGN.

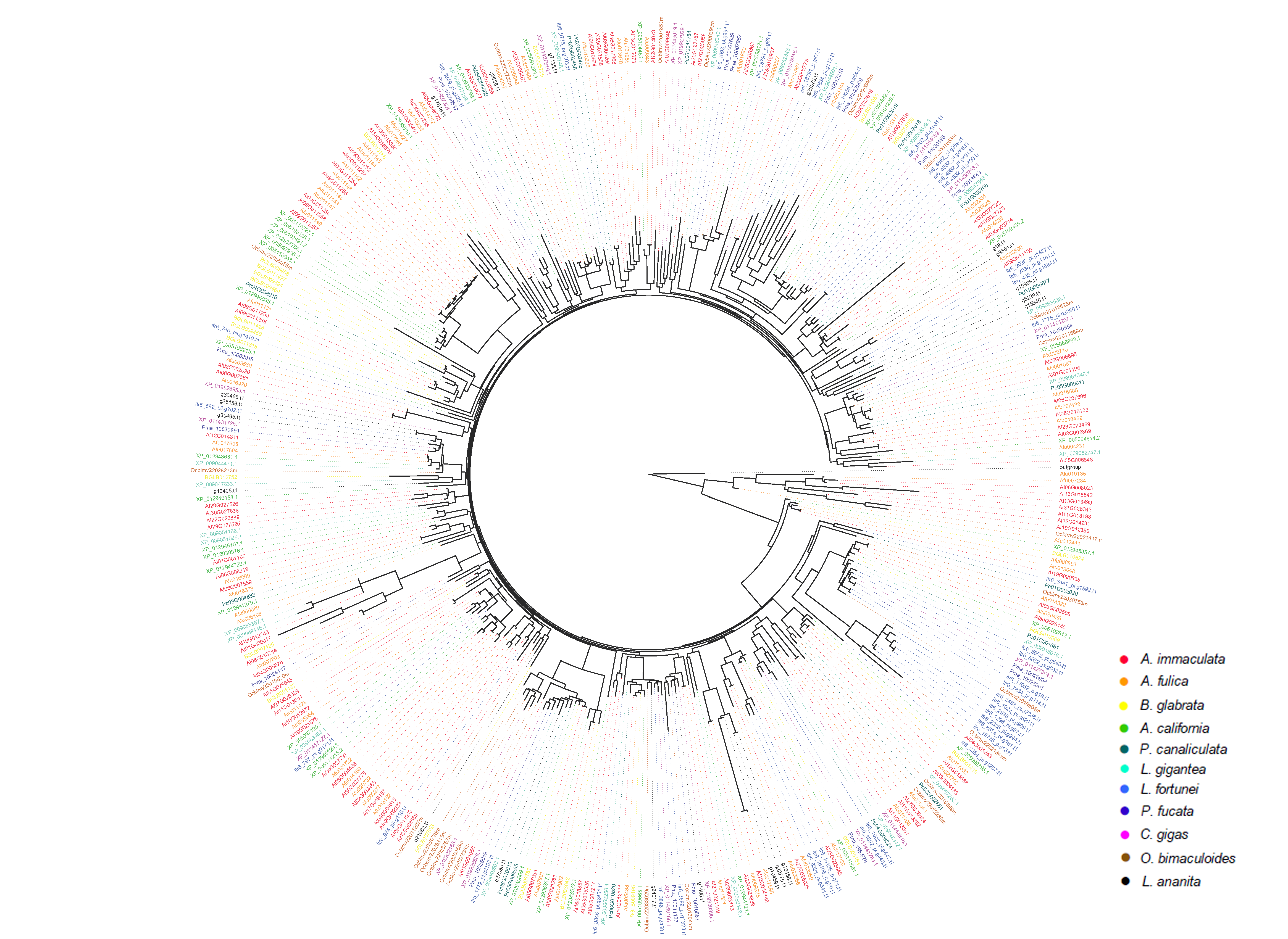

**Supplementary Fig. 17. Phylogenetic analysis of mucin in mollusc.** Phylogenetic analyses were performed using MEGA7 (Tamura et al., 2011) through maximum likelihood. The mucin genes from different species were indicated in circus filled with different colors. The mucin genes are highly expanded in *A. immaculata* and *A. fulica*, compared to other mollusc species.
